# CASP3-GSDME-mediated trophoblast pyroptosis contributes to systemic inflammation

**DOI:** 10.1101/2025.08.21.671642

**Authors:** Baoying Huang, Shilei Bi, Weinan Deng, Lili Du, Lijun Huang, Yifan Wang, Lizi Zhang, Zhoushan Feng, Tengfei Liu, Julia Kzhyshkowska, Zhaowei Tu, Haibin Wang, Jingsi Chen, Dunjin Chen, Shuang Zhang

## Abstract

Early-onset preeclampsia (EOPE) is associated with excessive apoptosis and inflammation, but the mechanistic link between these processes remains enigma. Here, we report elevated circulating pro-apoptotic proteins in EOPE patients at early pregnancy, along with concurrent CASP3 activation and GSDME cleavage in a subset of EOPE placentas. Using multiple trophoblast cell lines, we demonstrate that trophoblast cells, which highly express GSDME, undergo a shift from apoptosis to CASP3-dependent pyroptosis, driving inflammation. Notably, pyroptotic trophoblasts further induce pro-inflammatory macrophage polarization within placental villi organoids, establishing a feedback loop that amplifies both trophoblast pyroptosis and inflammatory responses in trophoblast organoids-macrophage assembloids. *In vivo*, CASP3-GSDME-mediated trophoblast pyroptosis contributes to systemic inflammation in wild-type pregnant mice but not in *Gsdme^-/-^* mice. Screening of EOPE prevention drugs reveals Vitamin D as a suppressor of GSDME activation and pyroptosis in trophoblast cells. These findings highlight the CASP3-GSDME axis as a promising therapeutic target for preeclampsia prevention.

## Introduction

Preeclampsia (PE) stands as a significant contributor to maternal and perinatal morbidity and mortality, afflicting 2 to 4 out of every 100 pregnancies worldwide ^1–4^. It manifests in two distinct forms delineated by gestational age at the time of diagnosis or delivery: early-onset PE (EOPE, < 34 weeks) and late-onset PE (LOPE, ≥ 34 weeks)^5^. Compared to late-onset counterpart, early-onset preeclampsia is considered to be associated with a greater incidence of the HELLP syndrome, abnormal uterine artery Doppler waveforms, atherosis, placenta lesions, small for gestational age, and fetal growth restriction ^6,7^. Presently, the sole remedy available for PE is the termination of pregnancy, a measure fraught with health risks for short and long-term consequences of both the mother and the fetus ^8–13^. The pathogenesis of EOPE is primarily attributed to placental dysfunction, wherein the placenta releases pro-inflammatory factors during the second stage, inciting a systemic inflammatory response that culminates in the clinical manifestations of PE ^14^. Nonetheless, a comprehensive elucidation of the mechanisms underlying how placental abnormalities trigger the ensuing inflammatory cascade in PE remains elusive.

In normal placental development, apoptosis plays a crucial role in cytotrophoblast fusion, syncytiotrophoblast turnover, and placental invasion ^15,16^. Apoptosis is stimulated by factors like hypoxia and oxidative stress in placentation, triggering both extrinsic and intrinsic pathways that ultimately activate caspases and release apoptotic bodies into the maternal circulation ^17^. Excessive apoptotic cell death in placentation has been observed in PE cases and is considered as one of the causes ^18,19^. Nevertheless, apoptosis does not trigger inflammation as apoptotic bodies has intact membrane and are phagocytosed ^20^, suggesting potential involvement of other cell death forms.

Pyroptosis, also known as inflammatory necrosis, is a caspase-dependent program, showing DNA damage and chromatin condensation ^21^. Unlike apoptosis, pyroptotic cells exhibit swelling and the appearance of bubble-like protrusions on the cellular membrane before eventual rupture ^22,23^. Gasdermin proteins are key executors involved in pyroptosis, and their cleavage fragments form pores in the plasma membrane, thereby releasing pro-inflammatory cytokines including IL-1β and IL-18, as well as HMGB1 and ATP ^24^. There are six members of the human gasdermin family, including GSDM A-D, GSDME (also known as DFNA5) and DFNB59 ^25^. Specific gasdermin involved in pyroptosis depends on the context, the activation of caspase-1 and caspase 11/4/5 cleave GSDMD and release the N-terminal domain that can oligomerize to form pores in the cell membrane, while Caspase 3 inactivate GSDMD mediated pyroptosis by cleaving it at Asp87 site ^26^. By contrast, GSDME is cleaved by active CASP3 at Asp270 to release a necrotic N-GSDME fragment to induce pyroptosis directly ^27^, alternatively, N-GSDME fragment in turn activates the NLRP3 inflammasome, leading to activation of the caspase 1/GSDMD cascade ^28,29^. GSDME mediated pyroptosis is increasingly reported to be involved in modulation of cancer progression and inflammatory diseases, while GSDME disruption skews pyroptotic death to apoptosis at sites of inflammation ^30–33^. A recent study showed that GSDME is highly expressed in trophoblast cells ^34^, indicating its potential role in mediating pyroptosis and subsequent inflammation in EOPE.

In this work, we demonstrate that GSDME, which is extensively expressed in trophoblast cells, switches CASP3 mediated apoptosis to pyroptosis, leading to a release of inflammatory cytokines. Remarkably, we elucidate a feedback loop between pyroptotic trophoblasts and pro-inflammatory macrophages within EOPE placentas. These results highlight CASP3 as a predictive marker and position the CASP3-GSDME pathway as a potential therapeutic target for preeclampsia prevention.

## Results

### High Relevance of CASP3 Activation and GSDME Cleavage in EOPE Placentas

To explore the potential interactions between death pathways in EOPE placentas, we conducted western blot analysis to examine key proteins involved in apoptosis (CASP3, Cleaved CASP3), necroptosis (p-RIPK3, RIPK3, p-MLKL, MLKL), and pyroptosis (GSDME-FL, GSDME-N) in placental samples from 14 EOPE patients and 12 gestational age-matched preterm deliveries (Supplementary Figure 1A). As shown in Figures 1A-B and Supplementary Figure 1B, we observed activation of the apoptotic pathway in EOPE placentas, marked by elevated levels of cleaved CASP3. Additionally, a significant increase in GSDME-N expression was noted in EOPE placentas. Notably, GSDME cleavage was observed concurrently with CASP3 activation in 7 out of 14 EOPE placenta (Figures 1A-C and Supplementary Figure 1B). Moreover, the key necroptotic markers, including total RIPK3, phosphorylated RIPK3, total MLKL, and phosphorylated MLKL, showed no notable alterations in EOPE placentas compared to age-matched control group (Figure 1A and Supplementary Figures 1C). Immunohistochemical staining of GSDME and cleaved CASP3 further confirmed elevated levels of these proteins in the villi of EOPE placentas, suggesting a potential role for CASP3-GSDME-mediated pyroptosis in EOPE pathology (Figures 1D-E).

**Figure 1.**
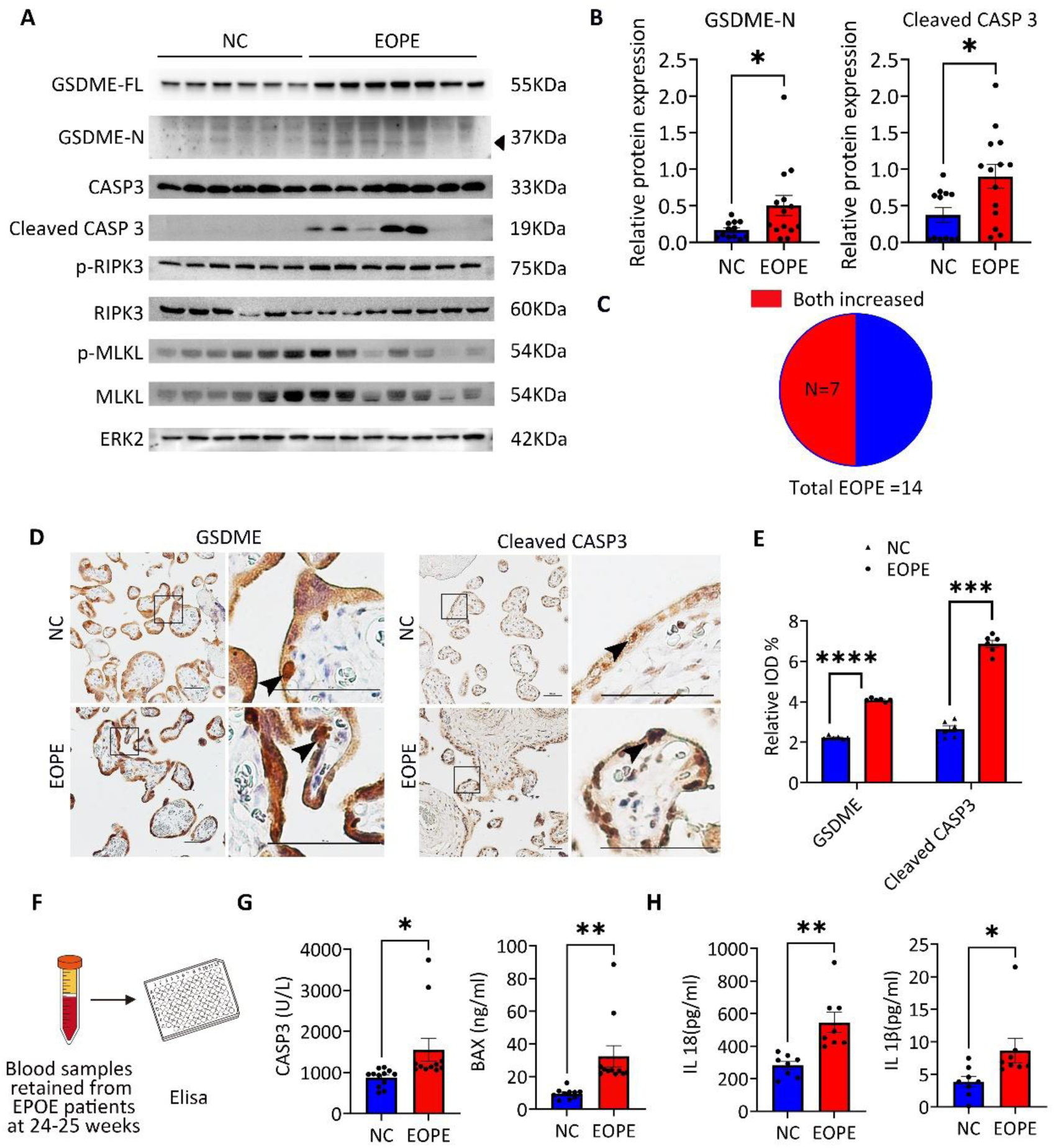
High Relevance of CASP3 Activation and GSDME Cleavage in a Subset of EOPE Placentas. (A) Immunoblots of pyroptosis-related proteins (GSDME-FL, GSDME-N, CASP3 and Cleaved CASP3) and necroptosis-related proteins (p-RIPK3, RIPK3, MLKL and p-MLKL) in lysate of placenta villi from NC and EOPE groups. ERK2 was used as a loading control. (B) Relative quantification of GSDME-N, Cleaved CASP3 in the normal controls(n=12) and EOPE(n=14) placentas by Western blot. Error bars, mean ± SEM. The data were analyzed by Student’s t-test, * p<0.05. (C) Pie chart showing the proportion of EOPE patients exhibiting increased levels of both GSDME-N and cleaved CASP3 in placental villous lysates. (D) Representative immunostaining pictures of GSDME and Cleaved CASP3 in the placenta of NC and EOPE group. The arrows indicated the positive trophoblast cells. Scale bars, 50 μm. (E) Relative quantification of GSDME and cleaved CASP3 immunostaining signals in placentas from NC and EOPE groups. Error bars, mean ± SEM. The data were analyzed by Student’s t-test, n ≥ 3, **** p<0.0001, *** p<0.001. (F) Schematic diagram of ELISA for detecting blood samples. (G) CASP3 expression in plasma samples from EOPE patients and matched controls at 24–25 weeks of gestation, as measured by ELISA. PE, n=11; NC, n=12. Error bars, mean ± SEM. The data were analyzed by Student’s t-test, * p<0.05. (H) IL-1β and IL-18 levels in plasma from patients with early-onset preeclampsia (EOPE) and matched controls at 24–25 weeks of gestation, as measured by ELISA. EOPE, n=8; NC, n=8. Error bars, mean ± SEM. The data were analyzed by Student’s t-test, * p<0.05.

Based on this temporal relationship, we postulated that circulating apoptotic proteins could represent early predictive biomarkers, as they appear before clinical symptom onset. We performed this OLINK proteomics in the blood samples of pregnant women between gestation weeks 12-16 who were subsequently diagnosed with EOPE (n=5) and control groups (n=4) (Supplementary Figure 1D). Among the proteins upregulated in the EOPE group, apoptotic markers such as BAX, CASP3, DECR1, and MAGED1, as well as the epithelial protein EPCAM, were of particular interest. These findings suggest that excessive apoptosis may serve as an early predictor of EOPE, potentially associated with impaired placental development (Supplementary Figure 1E). We further quantified BAX and CASP3 levels by ELISA in blood samples obtained from EOPE patients at 24–25 weeks of gestation (Figure 1F). Compared to the control group, these patients exhibited significantly elevated levels of BAX and CASP3 in plasma (Figure 1G).

Previous studies have shown that GSDME can exacerbate inflammation by mediating the release of cytokines such as interleukin-1β (IL-1β) and IL-18 ^35,36^, both of which have been implicated in EOPE pathology. To investigate whether CASP3-GSDME-mediated pyroptosis contributes to the inflammatory response in EOPE, we measured the levels of IL-1β and IL-18 in plasma samples from EOPE patients (n=6). Elevated concentrations of both cytokines were observed compared to controls (Figure 1H). These findings suggest that the apoptosis-pyroptosis transition plays a critical role in driving inflammation in EOPE placentas.

### Excessive Apoptosis induce GSDME-mediate pyroptosis in trophoblast cells

To confirm the above hypothesis, we utilized an apoptosis inducer, TNFα and SM164 (T/S hereafter), to trigger apoptosis in trophoblast cell lines including human trophoblast stem cell (hTSC), BeWo cells and HTR-8/SVneo cells (Figure 2A)^37,38^. Trophoblast cells underwent apoptosis under the condition of T/S, exhibiting cell shrinkage and fragmentation into apoptotic bodies (Figure 2A). We then conducted immunoblot on pyroptosis related proteins on trophoblast cell lines treated with different concentrations of T/S. Our results showed that cleaved CASP3 and GSDME-N increased along with higher T/S concentration in all trophoblast cell lines including hTSC, BeWo cells and HTR-8/SVneo cells (Figure 2C-E). These findings were consistent with the results observed in EOPE patients (Figure 1A-C). High Mobility Group Box 1 (HMGB1), a Damage-Associated Molecular Patterns (DAMP) molecule, is only released under conditions of cell lysis and serve as an indicator of pyroptosis ^40,41^. The supernatant was collected 24 hours after T/S treatment and the protein level of HMGB1 was detected by western blot. As shown in Figure 2F-H, HMGB1 levels were significantly elevated at TNFα concentrations exceeding 20 ng/ml. Similarly, lactate dehydrogenase (LDH), an enzyme that can be detected in the process of pyroptosis when the cell plasma membrane breaks ^42^, was progressively elevated with increasing TNFα concentrations in all trophoblast cells (Figure 2I). Notably, to further validate our findings, we conducted the aforementioned experiments using primary cytotrophoblasts, and the results consistently aligned with those obtained from other cell lines (Supplementary Figure 2). This reinforces the reliability of our data and conclusions.

**Figure 2.**
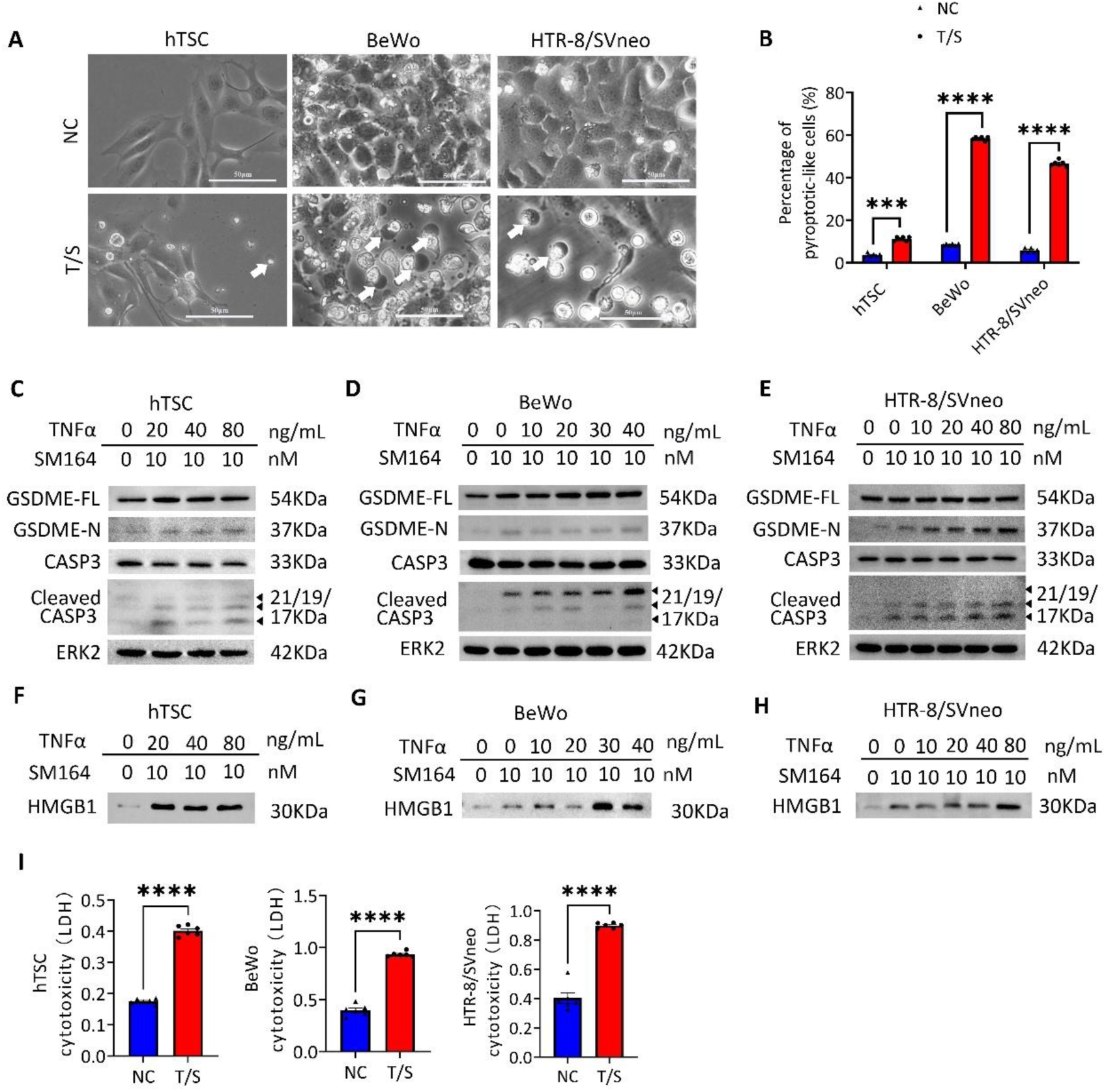
GSDME-mediate pyroptosis was induced upon activation of apoptosis in trophoblast cells. (A)Phase-contrast images of hTSC, BeWo cells and HTR-8/SVneo cells treated with SM164 and TNFα (T/S here after) after 24 hours. White arrows indicated the pyroptotic-like cells. Scale bars, 50 μm. (B) Percentage of pyroptotic-like cells following T/S treatment. Data are representative of three independent experiments. Error bars, mean ± SEM. The data were analyzed by Student’s t-test, n ≥ 3, **** p<0.0001, *** p<0.001. (C-E) Immunoblots of GSDME-FL, GSDME-N, CASP3 and Cleaved CASP3 in hTSC, BeWo cells and HTR-8/SVneo cells treated with SM164 and different TNFα concentrations (0, 10, 20, 40, 80ng/ml) after 24 hours. ERK2 was used as a loading control. (F-H) Immunoblots of HMGB1 in the supernatant of hTSC, BeWo cells and HTR-8/SVneo cells after treatment with T/S. (I) Cytotoxicity assay in hTSC, BeWo and HTR-8/SVneo cell after T/S treatment based on the detection of released LDH. The experiments were performed in triplicates. Error bars, mean ± SEM. The data were analyzed by Student’s t-test, n ≥ 3, ****p<0.0001.

In primary cultured trophoblasts treated with brefeldin A (BFA, an ER stress inducer), cobalt chloride (CoCl₂, a hypoxia mimic), or T/S (inducing excessive apoptosis), each representing different pathological conditions for preeclampsia, we observed pyroptosis-like cellular morphology as shown in Supplementary Figure 2A-2F. Western blot analysis further demonstrated that all three treatments effectively induced the cleavage of GSDME-N. Additionally, GSDMD cleavage was observed in the BFA and CoCl₂ treated groups, consistent with a previous study ^43,44^. These findings suggest that pyroptosis can be activated through distinct molecular pathways under varying conditions.

### Inhibition of GSDME or CASP3 reduces pyroptosis-like phenotype

To determine whether GSDME is the key director of pyroptosis in trophoblast cells, we used shRNA to knockdown the expression of GSDME in trophoblast cells. The expression of GSDME was efficiently decreased with sh*GSDME* infection in HTR-8/SVneo cells and BeWo cells as detected by western blot (Figure 3A and Supplementary Figure 3A). Immunoblot analysis further demonstrated significantly reduced GSDME expression and N-terminal cleavage in the sh*GSDME* group compared to sh*CTRL* following T/S treatment, despite robust CASP3 activation in both groups (Figure 3B). Consistent with reduced GSDME cleavage, HMGB1 levels in the supernatant of the sh*GSDME* group were significantly lower after T/S treatment, compared to the sh*CTRL* groups (Figure 3B). Interestingly, we observed the pyroptotic cells were significantly reduced *in* sh*GSDME* trophoblast cells upon T/S treatment when compared to the sh*control* (sh*CTRL*) group (Figure 3C and Supplementary Figures 3B-C). Additionally, total cell death was reduced, primarily due to a decrease in pyroptosis (Figure 3D). The release of LDH was also decreased with GSDME knockdown in both HTR-8/SVneo cells and BeWo cells (Figure 3E and Supplementary Figure 3D).

**Figure 3.**
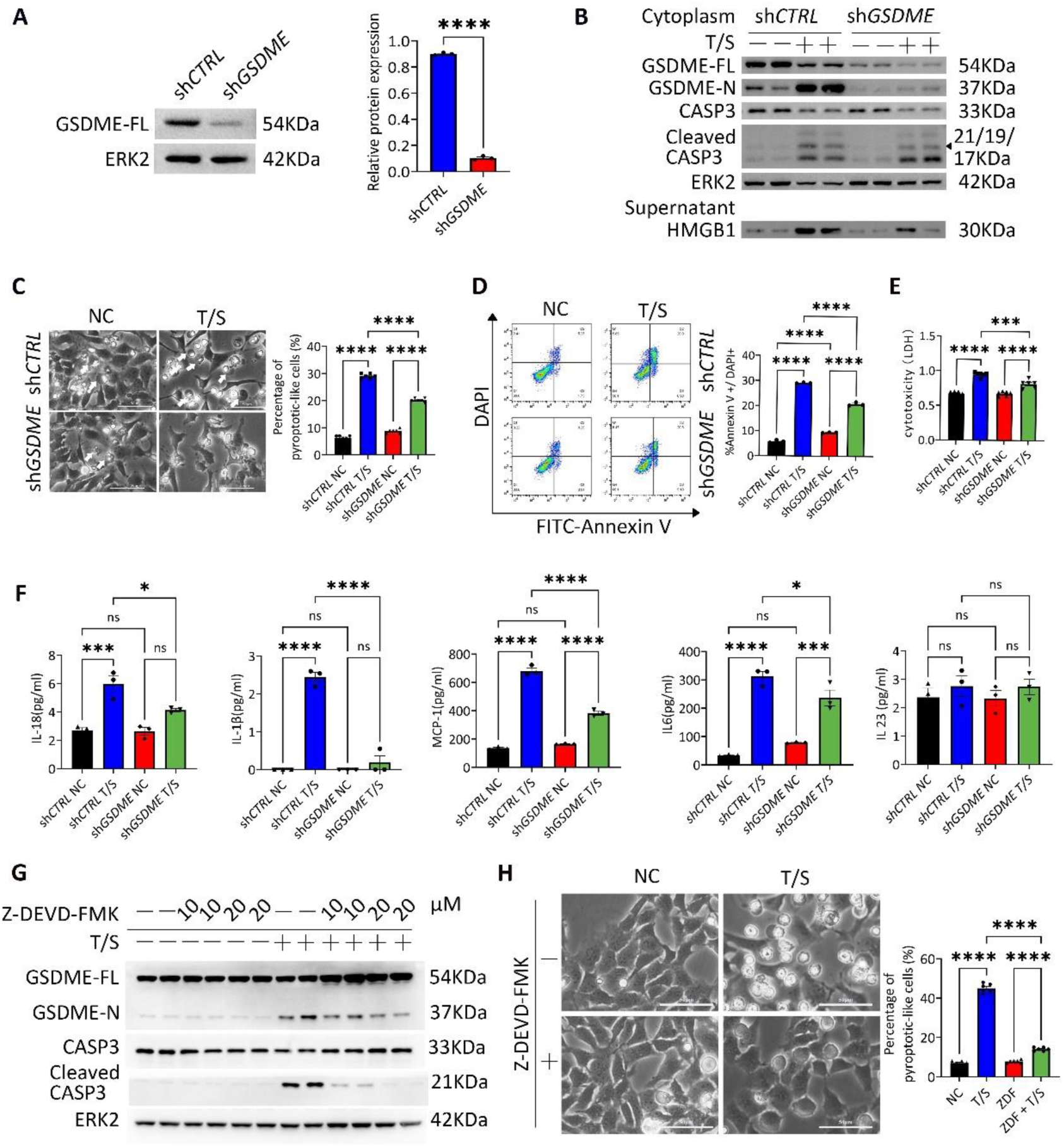
Inhibition of GSDME or CASP3 reduced pyroptosis-like phenotype and inflammation in trophoblast cells. (A) Immunoblots of GSDME and its relative quantification in sh*CTRL* and sh*GSDME* in HTR- 8/SVneo cells. Error bars, mean ± SEM. The data were analyzed by Student’s t-test, n ≥ 3, ****P < 0.0001. (B) Immunoblots of GSDME-FL, GSDME-N, CASP3, Cleaved CASP3 and HMGB1 in HTR-8/SVneo cells (sh*CTRL* and sh*GSDME*) treated with T/S. ERK2 was used as a loading control. (C) Phase-contrast images of sh*CTRL* and sh*GSDME* HTR-8/SVneo cells at 24 hours after T/S treatment. Arrows indicated the pyroptotic-like cells. Scale bars, 50 μm. Percentages of pyroptotic-like cells after T/S treatment. Error bars, mean ± SEM, n ≥ 3. The data were analyzed with a one-way ANOVA, ****p<0.0001. (D) Flow cytometry of Annexin V and DAPI in sh*CTRL* and sh*GSDME* HTR-8/SVneo cells treated with T/S. Percentages of Annexin V and DAPI double positive cells. Error bars, mean ± SEM, n ≥ 3. The data were analyzed with a one-way ANOVA, ****p<0.0001. (E) Comparison of LDH release of HTR-8/SVneo cells (sh*CTRL* and sh*GSDME*) treated with T/S after 24 hours. Error bars, mean ± SEM, n ≥ 3. The data were analyzed with a one-way ANOVA, **** p<0.0001, *** p<0.001. (F) Relative levels of IL-1β, IL-18, MCP-1, IL-6 and IL-23 in the supernatant of cultured sh*CTRL* and sh*GSDME* HTR-8/SVneo cells using the LEGENDplex™ inflammation panel. Error bars, mean ± SEM, n ≥ 3. The data were analyzed with a one-way ANOVA, * p<0.05, *** p<0.001, **** p<0.0001, ns, not significant. (G) Immunoblots of GSDME-FL, GSDME-N, CASP3 and Cleaved CASP3 in HTR-8/SVneo cells treated with T/S and Z-DEVD-FMK. ERK2 was used as a loading control. (H) Phase-contrast images of HTR-8/SVneo cells at 24 hours after T/S or Z-DEVD-FMK treatment. Arrows indicated the pyroptotic cells. Scale bars, 50 μm. Percentages of pyroptotic cells after T/S treatment. Error bars, mean ± SEM, n ≥ 3. The data were analyzed with a one-way ANOVA, ****p<0.0001, ***p<0.001.

It had been reported that GSDMD and GSDME perpetuate inflammation by mediating the release of cytokines such as IL-1β and IL-18 ^35,36^. As demonstrated in Figure 3F, T/S treatment significantly enhanced the secretion of both IL-1β and IL-18. Notably, GSDME knockdown partially attenuated this inflammatory response, as evidenced by reduced expression levels of these cytokines following T/S stimulation.

We next introduced the CASP3 inhibitor, Z-DEVD-FMK ^45,46^, to evaluate whether GSDME-mediated pyroptosis is CASP3 dependent. As shown in Figure 3G, immunoblot analysis confirmed that both the N-terminal fragment of GSDME and cleaved CASP3 were significantly decreased in the Z-DEVD-FMK-treated group compared to the control group after T/S treatment. Consistently, we observed a significant reduction in pyroptotic-like cells in Z-DEVD-FMK-treated trophoblast cells following T/S treatment compared to the control group. (Figure 3H). These findings suggest that reducing CASP3 activation or GSDME cleavage can alleviate pyroptosis.

### Pyroptotic trophoblasts promote pro-inflammatory macrophage polarization within placenta villi organoids

Fetal-origin Hofbauer macrophages have recently been identified as key mediators of inflammation, particularly in EOPE, but not in LOPE ^47,48^. To explore the impact of excessive apoptosis on placenta microenvironment during early pregnancy, we isolated placenta villi and cultured them as organoids (PVOs), which well retained both trophoblast cells and immune cells, following the protocol described in our previous publication ^49^ (Figure 4A). Morphologically, the PVOs exhibited a close resemblance to native placental villi after one week of *in vitro* culture (Figure 4B). The PVOs treated with T/S showed increased CASP3 activation and GSDME cleavage (Figure 4C), as well as elevated levels of inflammatory cytokines such as IL-1β and IL-18 in the supernatant (Figure 4D), in line with the observations in other trophoblast cells. Interestingly, we observed the macrophages within the PVOs treated with T/S displayed an inflammatory phenotype, showing decreased CD163, the classical M2 macrophage marker, and increased iNOS, a M1 macrophage marker, reflecting the imbalance of M1 iNOS+/CD163+ observed in PVOs of EOPE patients (Figure 4E).

**Figure 4.**
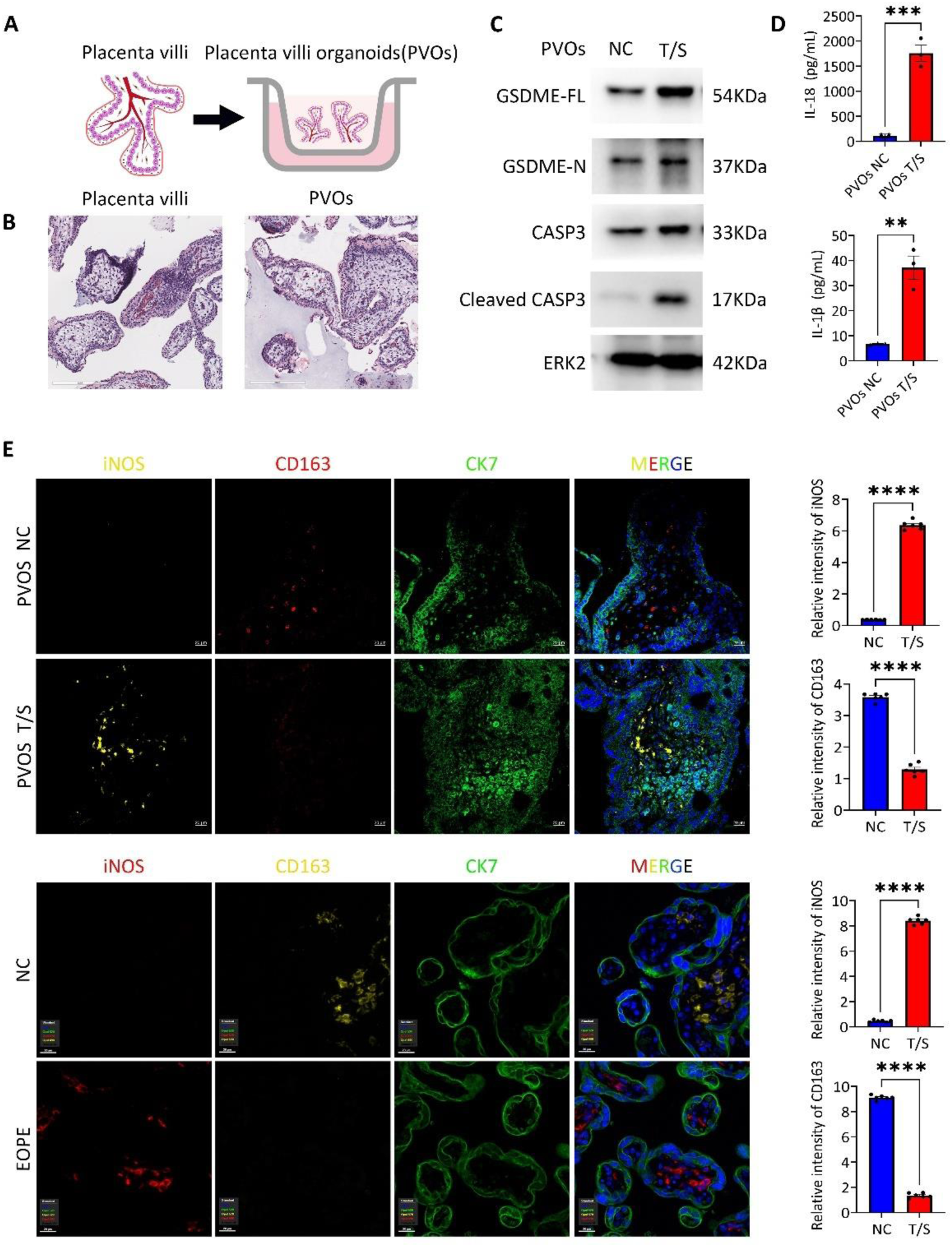
Pyroptotic trophoblasts drive pro-inflammatory macrophage polarization within placenta villi organoids. (A) Schematic representation of placenta villi organoids (PVOs) construction. (B) Haematoxylin and eosin (H&E) staining of placenta villi and PVOs. (C) Immunoblots of pyroptosis-related proteins (GSDME-FL, GSDME-N, CASP3 and Cleaved CASP3) in lysate of PVOs treated T/S after 24 hours. ERK2 was used as a loading control. (D) IL-18 and IL-1β level in the culture supernatant of PVOs treated T/S after 24 hours using the ELISA kit. Error bars, mean ± SEM. The data were analyzed by Student’s t-test, n ≥ 3, *** p<0.001, ** p<0.01. (E) Representative Immunofluorescence staining images of iNOS, CD163, CK7 and DAPI of PVOs treated T/S after 24 hours. Representative Immunofluorescence staining images of iNOS, CD163, CK7 and DAPI in the NC and EOPE placentas. The quantification of relative level of iNOS and CD163 in the PVOs treated T/S after 24 hours. Error bars, mean ± SEM. The data were analyzed by Student’s t-test, n ≥ 3, **** p<0.0001. The quantification of relative level of iNOS and CD163 in the NC and EOPE placentas. Error bars, mean ± SEM. The data were analyzed by Student’s t-test, n ≥ 3, **** p<0.0001.

### Pyroptotic trophoblasts and pro-inflammatory macrophages form a feedback loop that accelerates inflammation

To examine the potential association between macrophage polarization and trophoblast pyroptosis, we employed human trophoblast organoids (TOs) derived from placenta villi. Using this model system, we systematically evaluated the response of TOs to varying concentrations of T/S treatment. As expected, cleaved CASP3 and GSDME-N levels increased with higher T/S concentrations in TOs (Figures 5A-B), consistent with observations in other trophoblast lines. Furthermore, LDH levels also elevated in parallel with T/S concentrations, further validating TOs as a suitable model for studying the effect of pyroptotic trophoblast cells on macrophage polarization (Figure 5C).

**Figure 5.**
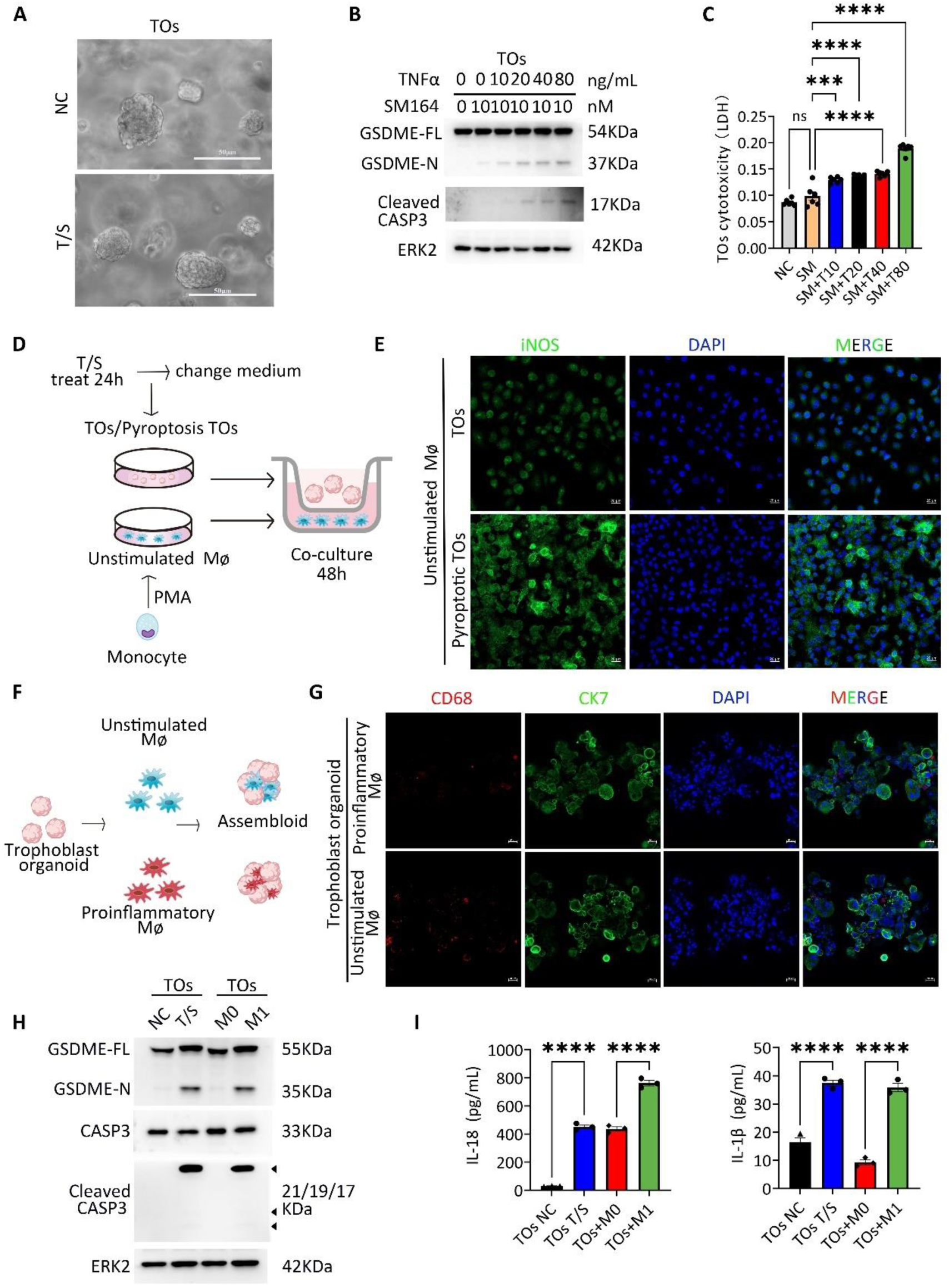
Pyroptotic trophoblasts and pro-inflammatory macrophages influence each other, promoting the inflammatory response. (A)Phase-contrast images of TOs treated with SM164 and TNFα (T/S here after) after 24 hours. Scale bars, 50 μm. (B) Immunoblots of GSDME-FL, GSDME-N and Cleaved CASP3 in TOs treated with SM164 and different TNFα concentrations (0, 10, 20, 40, 80ng/ml) after 24 hours. ERK2 was used as a loading control. (C) Cytotoxicity assay in TOs cell after T/S treatment based on the detection of released LDH. Error bars, mean ± SEM, n ≥ 3. The data were analyzed with a one-way ANOVA, ****p<0.0001, **p<0.01, *p<0.05. (D) Schematic representation of TOs and macrophage (Mφ) co-culture model construction. (E) Representative Immunofluorescence staining images of iNOS after co-culture of macrophages in the basal state with pyroptotic and non-pyroptotic TOs. (F) Schematic representation of TOs and Mφ assembloid construction. (G) Composite z stack confocal images of TOs and Mφ assembloid at day 3 after reaggregation containing unstimulated or proinflammatory macrophages stained with the antibodies against CD68, CK7 and DAPI. Scale bars, 20 μm. (H) Immunoblots of pyroptosis-related proteins (GSDME-FL, GSDME-N, CASP3 and Cleaved CASP3) in lysate of TOs treated T/S after 24 hours and TOs and Mφ assembloid containing unstimulated or proinflammatory macrophages. ERK2 was used as a loading control. (I) IL-18 and IL-1β level in the culture supernatant of TOs treated T/S after 24 hours and TOs and Mφ assembloid containing unstimulated or proinflammatory macrophages using the ELISA kit. Error bars, mean ± SEM, n ≥ 3. The data were analyzed with a one-way ANOVA. **** p<0.0001.

The interaction between macrophages and trophoblast cells was then evaluated through establishing a co-culture model of macrophage and TOs (Figure 5D). THP-1 cells were subjected to a 48-hour treatment regimen involving PMA, thereby facilitating the acquisition of unstimulated macrophage. TOs were treated with T/S for 24 hours, after which the medium was replaced, and co-cultures of TOs and unstimulated macrophage were maintained for 48 hours using transwell-polycarbonate inserts. Immunofluorescence staining revealed in the TOs T/S treatment group, the expression of pro-inflammatory macrophages marker iNOS was significantly increased (Figure 5D-E), indicating that pyroptotic trophoblasts promote pro-inflammatory macrophage polarization.

To investigate the influence of pro-inflammatory and unstimulated macrophages on trophoblast cells, we constructed an assembloid model combining macrophage and TOs (Figure 5F-G). Initially, THP-1 cells were subjected to a 48-hour treatment regimen involving PMA alone as well as PMA in conjunction with IFNγ, thereby facilitating the acquisition of unstimulated macrophage and pro-inflammatory macrophages. Pro-inflammatory macrophage induced increased expression of GSDME-N and cleaved CASP3 in TOs, suggesting their role in driving pyroptosis in trophoblasts (Figure 5H).

Furthermore, we measured IL-18 and IL-1β levels in the culture supernatants of assembloid and TOs treated T/S after 24 hours using ELISA. Both the TOs T/S treatment group and pro-inflammatory macrophage assembloid group showed significantly increased levels of IL18 and IL1β (Figure 5I), which were consistent with findings in EOPE placenta (Figure 1H) and PVOs (Figure 4D). These results suggest that pro-inflammatory macrophage polarization drive trophoblasts pyroptosis. Taken together, our findings highlight the crosstalk between pyroptotic trophoblasts and pro-inflammatory macrophage within the placenta villi, creating a feedback loop that exacerbates trophoblast pyroptosis and amplifies inflammatory cascades.

### CASP3-GSDME mediated placenta pyroptosis contributes to systemic inflammation*in vivo*

To explore whether local placental inflammation induced by GSDME-mediated trophoblast pyroptosis and its crosstalk with pro-inflammatory macrophages contributes to systemic inflammation *in vivo*, we simulated the heightened apoptosis observed in EOPE, by administering T/S to pregnant mice at embryonic days 8.5 (E8.5), E9.5 and E10.5. Age-matched control mice received injections of 0.9% NaCl. Placenta and fetuses were collected at E11.5 to evaluate the impact of T/S treatment (Figure 6A). Initially, we tested various doses of TNFα ranging from 0.5-2 μg/kg and analyzed placental samples via immunoblotting. As shown in Figure 6B, a dose-dependent increase in CASP3 activation and GSDME cleavage were observed, in line with our *in vitro* findings. We selected TNFα at 1 μg/kg for subsequent investigations, as this concentration initiated the observed phenotype. Consistent with our *in vitro* experiments, T/S treatment significantly enhanced CASP3 activation and GSDME cleavage in mouse placenta, the pro-inflammatory macrophage CD68 was upregulated and anti-inflammatory macrophage marker CD206 was downregulated (Figure 6C-E).

**Figure 6.**
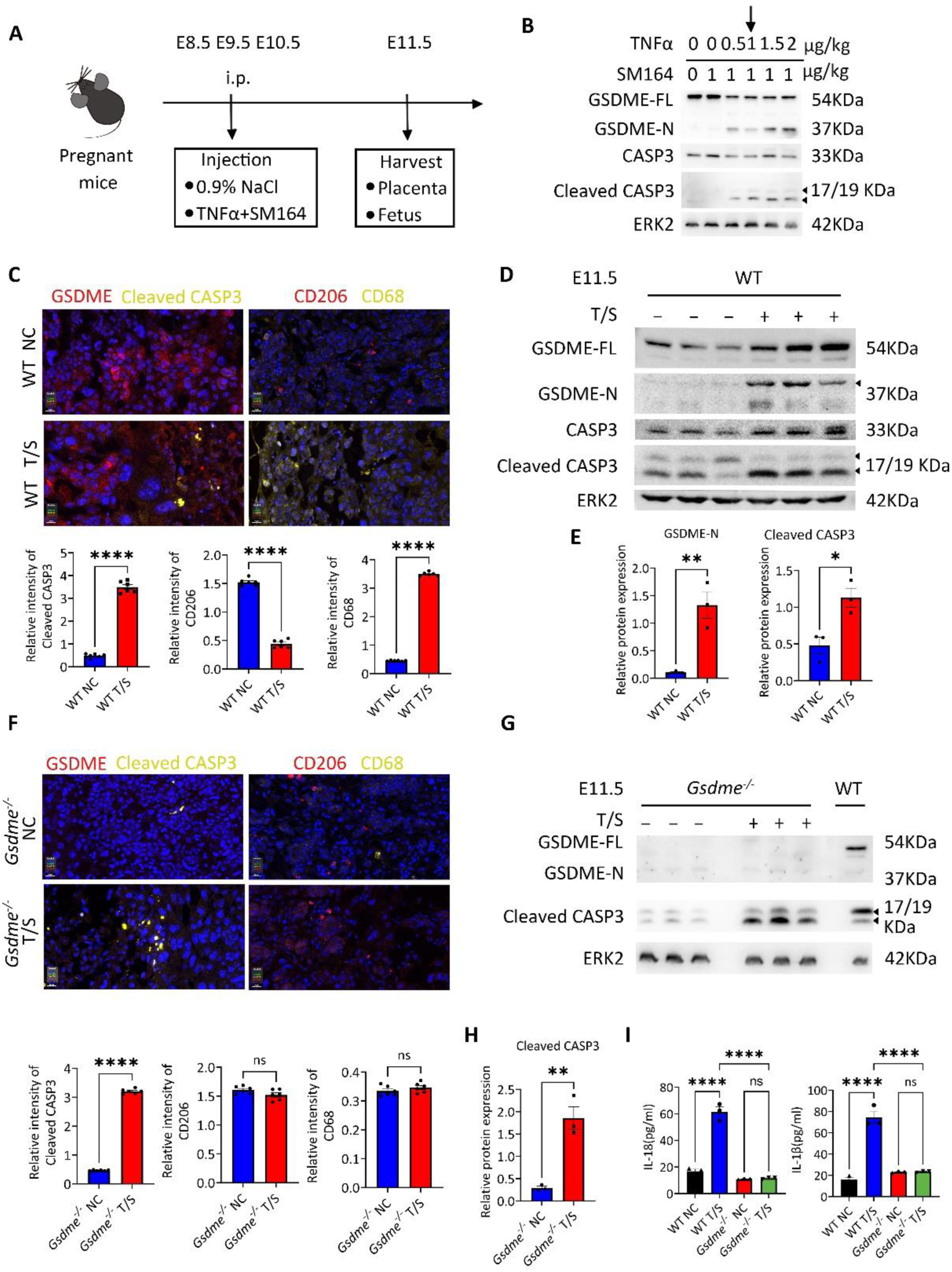
Caspase-3-GSDME-mediated placental pyroptosis plays a role in promoting systemic inflammation *in vivo*. (A) Scheme of the T/S treatment protocol in mice. The wild type (WT) and *Gsdme^-/-^* female mice were mated with WT male and *GSDME^-/-^*male mice respectively, and the pregnant mice were injected with TNFα and SM164 or with an equal volume of 0.9 % NaCl at embryonic day 8.5 (E8.5), E9.5 and E10.5. The mice were sacrificed at E11.5 and the placentas and fetues were collected for following experiments. (B) Immunoblots of GSDME-FL, GSDME-N, CASP3 and Cleaved CASP3 in placentas of WT mice treated with TNFα plus SM164. The dose of TNFα 1μg/kg and SM164 1μg/kg was used for the following injection. ERK2 was used as a loading control. (C) Representative immunofluorescence staining of GSDME, cleaved CASP3, CD206, and CD68 in placental tissue from T/S-treated mice. The quantification of relative level of Cleaved CASP3, CD206 and CD68 in placental tissue from T/S-treated mice. Error bars, mean ± SEM. The data were analyzed by Student’s t-test, n ≥ 3, **** p<0.0001. (D) Immunoblots of GSDME-FL, GSDME-N, CASP3 and Cleaved CASP3 in placental tissue from T/S-treated mice. ERK2 as the loading control. (E) Relative quantification of GSDME-N and cleaved CASP3 in placental tissue from T/S-treated mice, as determined by Western blot. Error bars, mean ± SEM. The data were analyzed by Student’s t-test, n ≥ 3, ** p<0.01, * p<0.05. (F) Representative Immunofluorescence staining images of GSDME, Cleaved CASP3 and CD206, CD68 in placental tissue from T/S-treated *Gsdme^-/-^* mice. Quantification of immunofluorescence signals for Cleaved CASP3, CD206 and CD68 in placental tissue from T/S-treated *Gsdme^-/-^* mice. Error bars, mean ± SEM. The data were analyzed by Student’s t-test, n ≥ 3, **** p<0.0001, ns, not significant. (G) Immunoblots of GSDME-FL, GSDME-N and Cleaved CASP3 in placental tissue from T/S-treated *Gsdme^-/-^* mice. ERK2 as the loading control. (H) Relative quantification of GSDME-N, Cleaved CASP3 in placental tissue from T/S-treated *Gsdme^-/-^* mice. Error bars, mean ± SEM. The data were analyzed by Student’s t-test, n ≥ 3, ** p<0.01. (I) IL-18 and IL-1β level in the serum of T/S-treated WT and T/S-treated *Gsdme^-/-^* mice using the ELISA kit. Error bars, mean ± SEM. The data were analyzed by Student’s t-test, n ≥ 3, **** p<0.0001, ns, not significant.

To ascertain whether local GSDME activation in the placenta contributes to systemic inflammation, we utilized a *Gsdme* knockout mouse model (*Gsdme^-/-^*). Following the same injection protocol as in wild-type mice (Figure 6A), immunofluorescence stain and western blot analysis of placenta revealed a complete absence of GSDME expression in *Gsdme^-/-^*placentas, confirming effective GSDME knockout (Figure 6F-G and Supplementary Figure 4). Furthermore, CASP3 activation was still upregulated following T/S treatment (Figure 6F-H).

Notably, we observed significantly elevated levels of the inflammatory cytokines IL-1β and IL-18 in the serum of T/S-treated mice compared to control mice, while this increase was attenuated in *Gsdme^-/-^*mice (Figure 6I), suggesting that GSDME-induced pyroptosis in mouse placenta contributes to systemic inflammation *in vivo*.

### Screening of EOPE Prevention Drugs Reveals Vitamin D Has a Role in Suppressing GSDME Activation and Trophoblast Cells Pyroptosis

Several medications are currently used in clinical practice for the prevention of EOPE. We examined whether commonly prescribed agents for EOPE prevention—GLP-1, Vitamin D, folic acid (FA), metformin (MET), and aspirin—achieve their therapeutic effects by suppressing GSDME activation (Figure 7A). As shown in Figure 7B, Vitamin D-treated trophoblast cells showed a decrease in GSDME cleavage when induced with T/S. However, we observed no evident reduction in pyroptotic cells and GSDME cleavage with GLP-1, FA, MET and aspirin treatments (Figure 7B-E). These findings suggest that Vitamin D may prevent the development of preeclampsia by suppressing trophoblast inflammation through the GSDME-CASP3 pathway.

**Figure 7.**
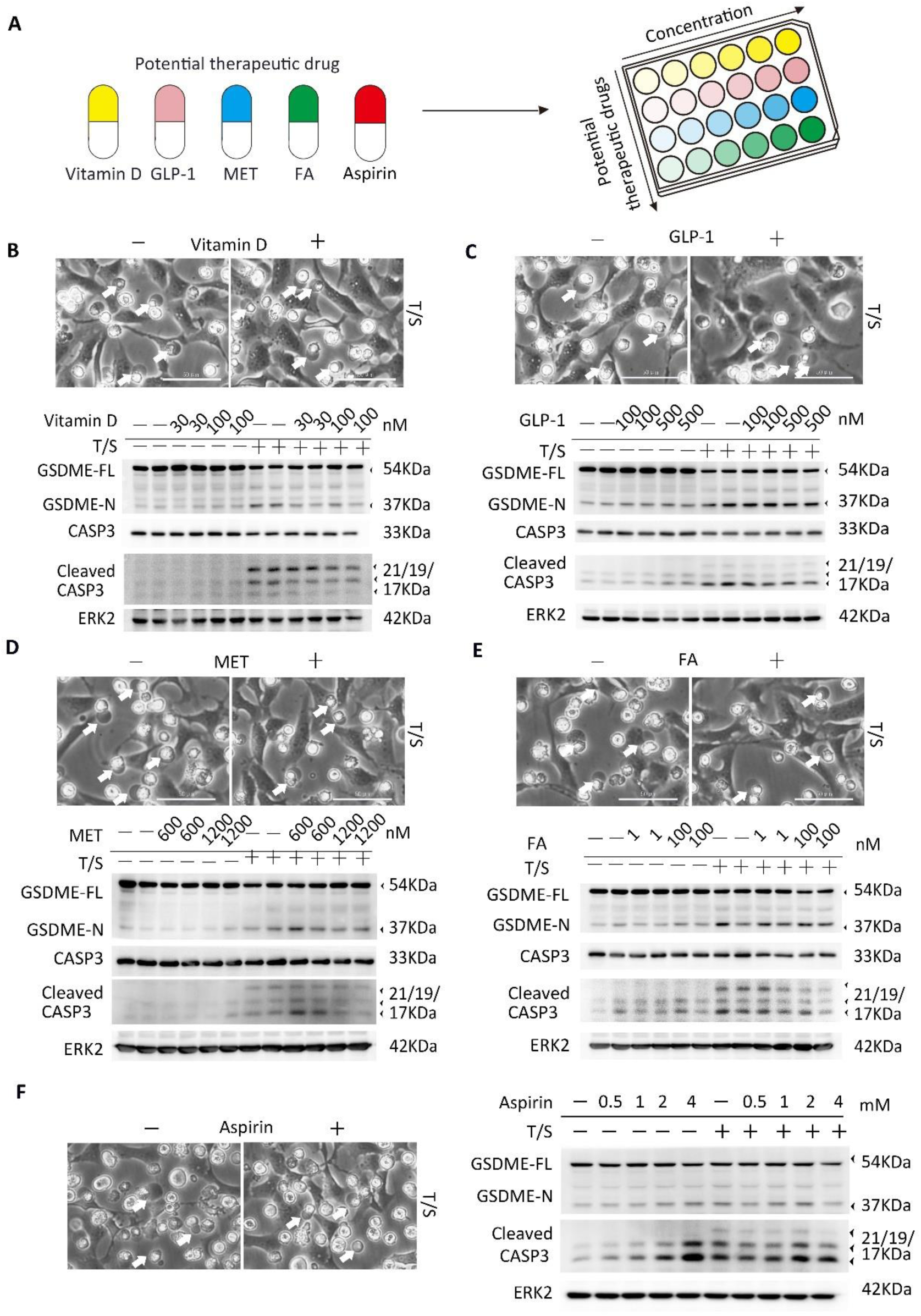
Screening the Effect of EOPE Prevention Drugs on Inhibiting Caspase-3-GSDME-Mediated Pyroptosis. (A) Scheme of the treatments of Vitamin D, GLP-1, MET and FA in HTR-8/SVneo cells. (B) Phase-contrast images, immunoblots of GSDME-FL, GSDME-N, CASP3 and Cleaved CASP3 of HTR-8/SVneo cells treated with Vitamin D and T/S after 24 hours. Arrows, the pyroptotic cells. Scale bars, 50 μm. (C) Phase-contrast images, immunoblots of GSDME-FL, GSDME-N, CASP3 and Cleaved CASP3 of HTR-8/SVneo cells treated with GLP-1 and T/S after 24 hours. Arrows, the pyroptotic cells. Scale bars, 50 μm. (D) Phase-contrast images, immunoblots of GSDME-FL, GSDME-N, CASP3 and Cleaved CASP3 of HTR-8/SVneo cells treated with MET and T/S after 24 hours. Arrows, the pyroptotic cells. Scale bars, 50 μm. (E) Phase-contrast images, immunoblots of GSDME-FL, GSDME-N, CASP3 and Cleaved CASP3 of HTR-8/SVneo cells treated with FA and T/S after 24 hours. Arrows, the pyroptotic cells. Scale bars, 50 μm. (F) Phase-contrast images, immunoblots of GSDME-FL, GSDME-N, CASP3 and Cleaved CASP3 of HTR-8/SVneo cells treated with Aspirin and T/S after 24 hours. Arrows, the pyroptotic cells. Scale bars, 50 μm.

## Discussion

We demonstrated that GSDME in trophoblast cells switched CASP3 mediated apoptosis to pyroptosis, leading to cell swelling and inflammation by releasing LDH, HMGB1 and inflammatory cytokines IL-18 and IL-1β. GSDME knockdown mitigated these adverse effects, and targeting activation of CASP3 by Z-DEVD-FMK reduced inflammation in trophoblast cell lines. These findings provide genetic evidence supporting the role of trophoblasts pyroptosis in mediating systemic inflammation of EOPE patients and highlight the potential of targeting the CASP3-GSDME pathway as a new prevention strategy for preeclampsia.

Our comparative analysis of placental samples from EOPE patients and age-matched pregnancies revealed a higher expression of the activated form of GSDME (N-GSDME) in EOPE placenta. Subsequent investigations on multiple trophoblast models indicated that CASP3 cleaves GSDME, converting trophoblast cell apoptosis to pyroptosis under conditions of consecutive apoptosis induction. This mechanism parallels observations in tumor cell death induced by chemotherapeutic agents ^28,29^. Notably, a previous study reported active Caspase-1, increased GSDMD cleavage, IL-1β, and IL-18 in EOPE placenta and primary human trophoblasts treated with BFA ^43^. However, our study demonstrates that GSDME can also be cleaved by BFA, as well as under hypoxic and apoptotic conditions, highlighting its critical role in mediating various pathological conditions. Additionally, we observed that the cleavage of GSDMD occurred concurrently with GSDME cleavage under BFA and CoCl₂ treatments, but not during T/S treatment. This suggests that pyroptosis may be activated through distinct molecular pathways depending on the specific pathological condition. Fetal-origin Hofbauer macrophages have recently been idelntified as key mediators of inflammation, particularly in EOPE, but not in LOPE ^47,48^. To replicate the conditions of excessive cytotrophoblast apoptosis observed in EOPE, we treated these PVOs isolated from 6-8 weeks pregnancies with an apoptosis inducer. Notably, this treatment led to an increase in M1 macrophages, resembling the inflammatory macrophage profile observed in placental villi during late pregnancy. These results hightlight excessive apoptosis as a primary trigger for the initiation of EOPE, aligning with previous studies^19^. Furthermore, co-culture experiments of macrophages with trophoblast spheroids (TSO) or assembloids comprising both cell types provided robust evidence that pyroptotic trophoblasts and pro-inflammatory macrophages establish a self-reinforcing inflammatory cascade within EOPE placentas.

Apart from pyroptosis, necroptosis represents another pro-inflammatory form of cell death, releasing intracellular damage-associated molecular patterns (DAMPs) that promote inflammation ^50^. Necroptotic morphology has been observed in placentas of mice exhibiting preeclampsia-like symptoms ^51^. In late preeclampsia, trophoblast cells undergo necroptotic cell death induced by ceramide ^18^. Key necroptosis effector molecules, including RIPK1 and SIRT2, have been detected in placental tissues of both normal pregnancies and those with PE ^52^. However, a recent study reported no evidence of necroptosis in placental tissue from EOPE patients ^43^, suggesting distinct inflammatory mechanisms between EOPE and LOPE. Consistent with this finding, we observed no significant alterations in necrosis, as evidenced by the minimal detection of p-RIPK3 and p-MLKL in EOPE placenta.

In conclusion, our study elucidates that CASP3 activation of GSDME-mediated pyroptosis in trophoblast cells at maternal-fetal interface. Pyroptotic trophoblast cells and the polarization of pro-inflammatory macrophages create a feedback loop, thereby promoting the trophoblast cell pyroptosis and inflammatory cascades. The evaluation of commonly used drugs and CASP3 inhibitors on trophoblast inflammation provides valuable insights for clinical therapeutic studies. CASP3 can serve as a predictive marker for the early prediction of EOPE (Figure 8).

**Figure 8.**
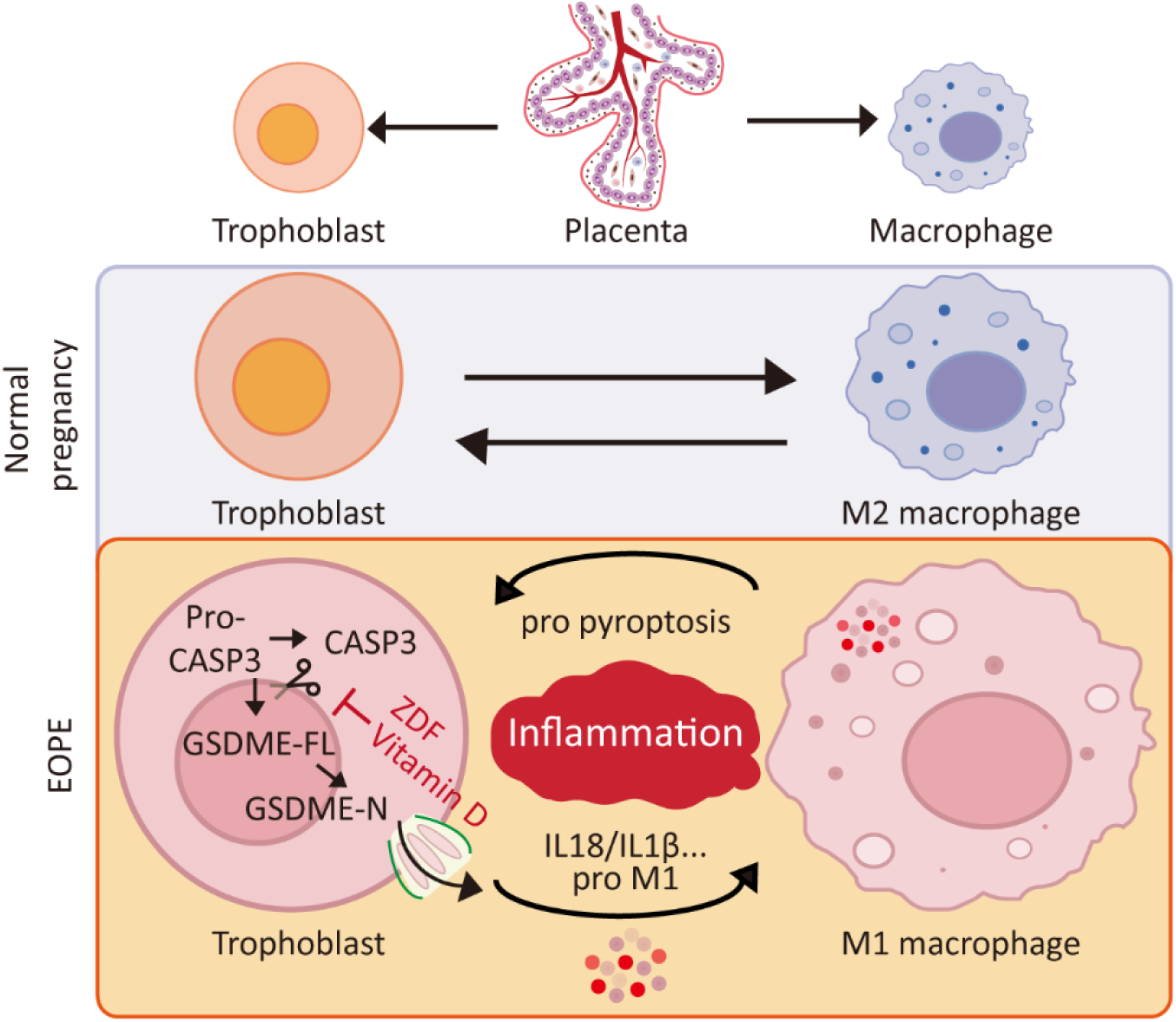
Schematic Representation of CASP3-GSDME-mediated trophoblast pyroptosis contributes to systemic inflammation. In trophoblast cells, treatment with the apoptotic inducer T/S (TNFα + SM164) induced a shift from CASP3-mediated apoptosis to pyroptosis, marked by cell swelling and inflammatory activation. This process was accompanied by the release of LDH, HMGB1, and the inflammatory cytokines IL-18 and IL-1β, which promoted the polarization of pro-inflammatory macrophages and further amplified pyroptosis in trophoblasts, thereby exacerbating systemic inflammation. Inhibition of CASP3 with Z-DEVD-FMK (ZDF) and Vitamin D attenuated inflammatory responses in trophoblast cell lines. T/S, TNFα plus SM164. ZDF, Z-DEVD-FMK.

## Materials and methods

### Human placentas tissues collection

Tissue samples used for this study were obtained with written informed consent from participants. The study was approved by the Medical Ethics Committee of the Third Affiliated Hospital of Guangzhou Medical University, Medical Research (No. 2018002). Placentas tissues at 28-34 weeks of gestation from EOPE patients or age-matched healthy pregnancies were collected from women who underwent caesarean deliveries from January 2017 to October 2022 at the Third Affiliated Hospital of Guangzhou Medical University, Guangdong, China. The diagnostic criteria for EOPE were established based on guidelines outlined by the International Society for the Study of Hypertension in Pregnancy ^56^. EOPE is characterized by the onset of hypertension occurring after 20 weeks of gestation, presenting with a systolic blood pressure of ≥140 mmHg and/or diastolic blood pressure of ≥90 mmHg, accompanied by proteinuria at 0.3 g/day. Alternatively, EOPE may manifest without proteinuria, but with the presence of one or more of the following symptoms: thrombocytopenia, liver function impairment, renal function impairment, pulmonary edema, neurological abnormalities, intrauterine growth restriction, or uteroplacental insufficiency. Normal pregnant controls do not exhibit pregnancy complications such as gestational or chronic hypertension, pregestational or gestational diabetes mellitus, or intrahepatic cholestasis of pregnancy. Collected tissues were snap-frozen in dry ice at the time of surgery and stored at −80 °C or liquid nitrogen. Part of the tissue was fixed in 4 % PFA for histopathological analysis. Detailed information of patients is listed in Supplementary Table 1.

### Blood Sample Collection

Blood samples used for the retrospective proteomic study of early-onset preeclampsia (EOPE) were obtained from the biobank of the Third Affiliated Hospital of Guangzhou Medical University. The samples were collected during routine clinical practice at 12–16 and 24–25 weeks of gestation from consenting patients who were subsequently diagnosed with EOPE at the same hospital between 2019 and 2020 (Detailed information is listed in Supplementary Table 1). Plasma was prepared from whole blood collected into tubes with EDTA by centrifugation at 3000 rpm for 15 mins at room temperature, after which the plasma was aliquoted and frozen for long-term storage at −80 °C until analysis.

### Blood Sample Proteome Profiling

The protein levels of all samples were measured in plasma using the Olink Explore 1536, which uses antibody-binding capabilities to detect the level a combination of 4 separate Olink Explore 384 panels, including Cardiometabolic panel, Inflammation panel, Neurology panel and Oncology panel. Data is presented as NPX (Normalized Protein eXpression) values, which is relative protein quantification unit on log2 scale.

### Animals

The animals were housed in the animal care facility of Guangzhou Medical University according to the guidelines for the care and use of laboratory animals. All experimental procedures were approved by the Animal Welfare Committee of Research Organization of Guangzhou Medical University. Two-month-old C57BL/6 wild-type mice were purchased from Guangdong Medical Laboratory Animal Center and used in the experiments. *Gsdme^-/-^* mice were obtained from Prof. Feng Shao laboratory at National Institute of Biological Sciences, Beijing, the mice were constructed as previously reported ^28^. The wild-type and *Gsdme^-/-^* female mice were mated with wild-type male and *Gsdme^-/-^* male mice respectively, and the pregnant mice were injected with TNFα (1 μg/kg) and SM164 (1 μg/kg) or with an equal volume of 0.9 % NaCl at embryonic day 8.5 (E8.5), E9.5 and E10.5 of pregnancy. The mice were sacrificed at E11.5 and the placentas, fetues and serum were collected for indicated experiments. The genotyping primers for *Gsdme^-/-^* mice are as follows: F1: 5′ −tCCATTACTGTGGCTAAAGAGGGGC −3′; R1: 5′−CTTCCTAAACTCCTGCGGAAGACAG −3′ R2: 5′−GGCCTCTAGCCCAGCATGC −3′.

### Cell culture

Primary trophoblast cells (PTCs) were isolated from placenta during the third trimester (37-40 weeks of gestation) as previously reported ^58^. Placentas were collected immediately after cesarean section from uncomplicated singleton pregnancies at term. Exclusion criteria included fetal anomalies, maternal pathologies, or placental abnormalities. Tissue was transported on ice and processed within 1 hour. Amniotic membranes were removed, and placenta was diced into 2 × 2 cm pieces. Tissue was washed with 0.9% NaCl (4–8 L) until the wash solution became clear. Vessels, chorionic and decidual plates, and blood clots were meticulously dissected to minimize contamination. 72 g of minced tissue was incubated in 300 mL pre-warmed (37 °C) 1 × HBSS (Thermo, Cat. 14185052) containing 720 U dispase II (Sigma, Cat. D4693) for 1 hour under agitation. DNase I (100 U/mL, Worthington, Cat. LS006344) was added for an additional 15 min to reduce clumping. The digest was sequentially filtered through 200 μm, 100 μm, and 40 μm sieves. The filtrate was centrifuged (500 × g, 20 min), and the pellet was resuspended in wash medium (DMEM/F12 + 1 % Penicillin-Streptomycin). A discontinuous Percoll gradient (70–5%, 3 mL layers) was overlaid with 6 mL of resuspended cells and centrifuged (1,200 × g, 20 min, no brake). The cytotrophoblast layer (density 1.048–1.062 g/mL, fractions 45–35%) was aspirated, washed, and treated with 0.05% trypsin (1 min, 37 °C, Gibco, Cat. 25300062) followed by DNase I (100 U/mL). Cells were filtered (40 μm), counted (trypan blue exclusion), and viability >90% was confirmed. Cells were plated at 4.0 × 10^5 cells/cm^2 in DMEM/F12 medium (Corning, Cat. 10-092-CV) containing 10 % fetal bovine serum (FBS, Lifeman, Cat.FBS001) and 1 % Penicillin-Streptomycin (Gibco, Cat. 15140122) under 21% O₂/5% CO₂ at 37 °C. Medium was changed daily.

Human trophoblast stem cells (hTSC) were isolated from placenta during the first trimester (6-8 weeks of gestation) as previously reported ^59^. hTSCs were cultured in hTSC medium supplemented with 0.1 mM 2-mercaptoethanol (Thermo Fisher Scientific, Cat. 21985023), 0.2 % FBS (Corning, Cat. 26219002), 0.5 % Penicillin-Streptomycin (Gibico, Cat. 15140122), 0.3 % BSA (Sigma, Cat. A9418), 1% ITS-X supplement (Wako, Cat. 094-06761), 1.5 mg/ml L-ascorbic acid (Wako, Cat. 013-12061), 50 ng/ml EGF (Wako, Cat. 053-07871), 2 mM CHIR99021 (Wako, Cat. 034-23103), 0.5 mM A83-01 (Wako, Cat. 035-24113), 1 mM SB431542 (Wako, Cat. 031-24291), 0.8 mM VPA (Wako, Cat. 227-01071) and 5 mM Y27632 (Wako, Cat. 036-24023).

The methodology for establishing trophoblast organoids (TOs) was adapted from our previously published article ^60^. The hTSC were harvested and re-suspended in ice-cold basic trophoblast organoid medium (bTOM) containing advanced DMEM/F12(Gibco, Cat.12634010) supplemented with 10mM HEPES (Gibco, Cat. 15630080), B27 (Gibco, Cat. 17504044), N2 (Gibco, Cat.17502048) and 2mM L-glutamine (Life Technologies, Cat. 25030-081). The cells were then centrifuged for 3 minutes at 1000rpm following which the cells were re-suspended in ice-cold advanced trophoblast organoid medium (aTOM) which is bTOM supplemented with 100ng/ml R-spondin (PeproTech, Cat.HZ-1328), 1μM A8301 (Wako, Cat. 035-24113), 100ng/ml recombinant human epidermal growth factor (rhEGF, Peprotech, Cat. AF-100-15), 50ng/ml recombinant human hepatocyte growth factor (rhHGF, PeproTech, Cat. 100-39), 2.5μM prostaglandin E2 (Sigma, Cat. P0409), 3μM CHIR99021 (Wako, Cat. 034-23103) and 100ng/ml Noggin (PeproTech, Cat. 120-10c). Growth factor reduced matrigel (Corning, Cat. 356231) was added to the aTOM cell suspension to reach a final concentration of 60%. 50μl of the cell solution containing 2.5x10^4^ cells was plated in the center of a 24-well plate. The solution rests as a dome-shaped droplet in the center of the well. The plates are then turned upside down and kept at 37°C for 15-30 minutes to ensure proper spreading of the cells in the solidifying matrigel domes. Finally, the plates are returned to their upward position and the domes are overlaid with 500μl of room temperature aTOM medium. The organoids are allowed to form for 10 days with fresh media being changed every 2 days.

The methodology for establishing placenta villi organoids (PVOs) was adapted from our previously published article ^49^. Collagen solutions were mixed as follows: Cellmatrix I-A (Wako, Cat. 637-00653), Advanced DMEM/F12 (Gibco, Cat. 12634010), 20mM HEPES (Thermo Fisher Scientific, Cat. 15630080) at a ratio of 8:1:1. The mixture was kept on ice to prevent gel formation until use. The formation of air bubbles should be avoided during mixing. To prepare the culture dish, Millicell culture plate inserts (Millipore, Cat. R1KB36634) with permeable membrane bottoms were placed in a 60 mm tissue culture dish. To create the bottom layer, 1 ml of the prepared reconstituted collagen solution was added to the inserts under sterile conditions. The culture dish with the inserts was incubated in an incubator at 37 °C for 30 min. Villi slices or villi were washed with 10 ml basal medium and resuspended ∼0.1 mg in 1 mL reconstituted collagen solution. The mixtures were layered on top of the pre-solidified bottom layer to form the double dish air-liquid culture system as described. 2 ml placenta villi organoid culture medium was added into the culture dish on the outer layer. The culture medium was changed twice a week. The basal medium consisted of Advanced DMEM/F12 (Gibco, Cat. 12634010), 10mM HEPES (Thermo Fisher Scientific, Cat. 15630080), 1X GlutaMAX (Life Technologies, Cat. 2268102), and 1% penicillin/streptomycin (Gibco, Cat. 15140-122). The placenta villi organoid culture medium consisted of the basal medium supplemented with 10 mM nicotinamide (Sigma, Cat. N0636), 1X B-27 (Invitrogen, Cat. 12587010), 1mM N-acetylcysteine (Sigma, Cat. A9165), 10μM SB431542 (Wako, Cat. 031-24291), 0.5mM A83-01 (Wako, Cat. 039-24111), 100 ng/mL EGF (Wako, Cat. 053-07871), 100 ng/ml FGF2 (Peprotech, Cat. 450-33), 50 ng/mL HGF (Peprotech, Cat. 100-39), 2μM Y-27632 (Wako, Cat. 030-24021), 2.5μM PGE2 (Sigma, Cat. P0409), and 50% (v/v) WRN-CM. WRN-conditioned medium was obtained from a commercially available ATCC® CRL-3276™ (ATCC) cell line.

HTR8/SVneo and BeWo cell lines were obtained from the American Type Culture Collection (ATCC, Manassas, VA, USA) authenticated using STR profiling by ATCC, and tested to be free from mycoplasma contamination. HTR8/SVneo cells were grown in 10 cm cell culture dish with 1640 medium (Corning, Cat. 10-040-CV) containing 10 % fetal bovine serum (FBS, Lifeman, Cat.FBS001) and 1% Penicillin-Streptomycin (Gibco, Cat. 15140122). BeWo cells were cultured in 10cm cell culture dish with DMEM/F12 medium (Corning, Cat. 10-092-CV) containing 10 % fetal bovine serum (FBS, Lifeman, Cat.FBS001) and 1 % Penicillin-Streptomycin (Gibco, Cat. 15140122). All cells were cultured at 37 ℃ in 5 % CO_2_ and the culture medium was replaced every two days. Static bright field cell images were captured using the Leica microscope (Leica, Cat. DMIL LED).

### THP-1 cells culture and differentiation

THP-1 cell lines were obtained from the American Type Culture Collection (ATCC, Manassas, VA, USA) authenticated using STR profiling by ATCC, and tested to be free from mycoplasma contamination. THP-1 cells were grown in 10 cm cell culture dish with 1640 medium (Corning, Cat. 10-040-CV) containing 10 % fetal bovine serum (FBS, Lifeman, Cat.FBS001), 0.05 mM 2-mercaptoethanol (Thermo Fisher Scientific, Cat. 21985023) and 1% Penicillin-Streptomycin (Gibco, Cat. 15140122). THP-1 cells (1× 10^6^/well) were seeded into 6-well plates in the presence of 100 ng/ml phorbol 12-myristate 13-acetate (PMA, MCE, Cat.HY-18739) and Interleukin 4 (IL 4, PeproTech, Cat. 200-04) or Interferon gamma (IFNγ, PeproTech, Cat. 300-02) for 48 h. Then, the medium was removed and THP-1 cells were cultured with fresh RPMI 1640 medium for an additional 24 h. For the indicated experiments, the PMA-treated THP-1 cells were co-cultured with TSO for up to 48 hours.

### Construction of TOs and macrophages assembloid

The TOs and macrophages assembloid were constructed with TSO and THP-1 cells-induced macrophages. Briefly, TOs were released with Cultrex Organoid Harvesting Solution (R&D, Cat.3700-100-01) at Day 10-12 and macrophages were dissociated with Accutase after day 3 of the differentiation procedure.

The TOs and macrophages were reaggregated with approximately 95-98% TOs and approximately 2-5% macrophages in advanced trophoblast organoid medium (aTOM) supplemented with 10% fetal bovine serum (FBS, Lifeman, Cat.FBS001) using low-attach U plates. 48 hours later, the cells self-assembled into organoids.

### Apoptosis and pyroptosis induction and block

Using TNFα (PeproTech, Cat. 300-01A) and SM164 (Apexbio, Cat. A8815) as an apoptosis inducer. Apoptotic response were introduced as follows: TNFαand SM164 at the indicated concentrations in each experiment. ER stressors inducer Brefeldin A (CST, Cat. 9972) weas added to Primary Trophoblast Cells (PTCs) for 24 hour and the concentrations (1, 2.5, 5 μg/ml). Cobalt Chloride (CoCl_2_, Sigma-Aldrich, Cat. C8661) was used to simulate hypoxia inPTCs and the concentrations (200μM). To block pyroptosis induction, inhibitors including Z-DEVD-FMK (APEXBIO, Cat. A1920), Beinaglutide (GLP-1, MedChemExpress, Cat. HY-P3463), Vitamin D (VD, Sigma, Cat. D1530), Folic Acid (FA, Sigma, Cat. F7876), Aspirin (Selleck, Cat. S3017) and Metformin hydrochloride (MET, MedChemExpress, Cat. HY-17471A) was added 1 hour before T/S treatment.

### Plasmids construction and shRNA knockdown

The shRNA of GSDME were designed by Genetic Perturbation Platform from BROAD institute and primers (F: 5’- CCGGGCTTCTAAGTCTGGTGACAAACTCGAGTTTGTCACCAGACTTAGAAGCTTTTTG-3’; R:5’-AATTCAAAAAGCTTCTAAGTCTGGTGACAAACTCGAGTTTGTCACCAGACTTAGAAGC-3’) were synthesized by Shanghai General Biological Engineering Co, LTD. The shRNAs were cloned into the pLKO.1-Puro lentiviral vector. To generate lentiviral vectors, HEK293FT cells were cotransfected with lentiviral vectors, psPAX2, and pMD2.G at a ratio of 10:7.5:2.5 using jetPRIME Transfection Reagent (polyplus, Cat.114-15), following the manufacturer’s instructions. Media were replaced 8 hours post-transfection, and viral supernatants were collected 72 hours later after passing through a 0.45µm filter. The collected supernatants were aliquoted and stored at -80°C. To achieve GSDME knockdown, the HTR8 and BEWO were infected with *shGSDME* lentivirus and the cells were selected by puromycin.

### Methanol/Chloroform Protein Precipitation

Method to process 100μL of protein sample, it can be scaled up or down. Add 400μL of Methanol to a sample of 100μL volume and vortex well. Add 100μL of Chloroform and vortex well. Add 300μL of ddH_2_0 (PS: sample should look cloudy) and vortex well.

Spin 2 min, 14,000 g and then pipette off the top aqueous layer (PS: do not lose protein - protein exists between layers and may be visible as a thin wafer). Add 400μL of Methanol and then vortex well, spin 3min, 14,000 g. Pipette as much Methanol as possible from the tube without disturbing the pellet. Speed-Vac to dryness but avoid drying too long as this makes the pellet harder to re-solubilize. Solubilize the pellet in buffer appropriate for downstream process.

### Immunoblots analysis

The cells were washed once with cold PBS, lysed with cell lysis buffer (Beyotime, Cat. P0013) on ice for 30 min and shake every 10 min, followed by centrifugation at 12,000×g for 30 min. The supernatant was collected and protein concentration were determined using a BCA Protein Assay kit (Thermo, Cat. 23228). Equal amount of protein was separated by sodium dodecylsulfate-polyacrylamide gradient gel electrophoresis, and transferred onto a PVDF membrane, followed by primary and secondary antibodies incubation. Immunoblots images were developed using the SageCapture gel imaging system (ChampChemi610, Beijing, China). The Image J was employed for quantitative analysis and the relative protein expression level was determined by using ERK2 as loading controls. Primary antibodies: CASP1/p20/p10 Polyclonal Antibody (Proteintech, Cat. 22915-1-AP, dilution ratio of 1:1000), Cleaved Gasdermin D (CST, Cat. 36425S, dilution ratio of 1:1000), GSDME-N-terminal (Abcam, Cat. ab215191, dilution ratio of 1:1000), HMGB1 antibody (Abcam, Cat. ab18256, dilution ratio of 1:1000), CASP3 antibody (Sigma, Cat. HPA002643, dilution ratio of 1:1000), Cleaved CASP3 antibody (CST, Cat. 9664S, dilution ratio of 1:1000), MLKL antibody (Abcam, Cat. ab194699, dilution ratio of 1:1000), p-MLKL (Abcam, Cat. ab196436, dilution ratio of 1:1000), p-RIPK3 (Abcam, Cat. ab209384, dilution ratio of 1:1000), RIPK3 (Santa, Cat. sc-374639, dilution ratio of 1:200), ERK2 (Santa, Cat. sc-1647, dilution ratio of 1:400).

### Immunostaining and immunofluorescence

All the samples from the biopsy were fixed in formalin and embedded in paraffin for section. The 5 μm tissue sections were deparaffinized and dehydrated, subjected to antigen retrieval by autoclaving in 10 mM sodium citrate solution at 120°C for 15 min. For IHC, endogenous peroxidase activity was quenched in 3% H_2_O_2_ in methanol for 10 minutes, and sections were blocked in 5% BSA in PBS for 1 hour. Primary antibodies against GSDME-N-terminal (Abcam, Cat. ab215191, 1:200), CASP3 antibody (Sigma, Cat. HPA002643, 1:200), Cleaved CASP3 antibody (CST, Cat. 9664S, 1:200), MLKL antibody (Abcam, Cat. ab194699, 1:200), RIPK3 (Santa, Cat. sc-374639, 1:200) were added overnight at 4°C, then slides were washed in PBS three times for 10 minutes, incubated with secondary antibodies for 1 hour at room temperature, and washed again. and incubated with the secondary antibody at room temperature for 1 hour. The signals were visualized by diaminobenzidine (DAB) staining (ZSGB-BIO, Cat. ZLI-9019) at room temperature. IHC slides were counterstained with hematoxylin and scanned using an Olympus microscope (Cat. BX43). For IF, after antigen retrieval, slides were washed in PBS three times for 10 minutes and then blocked in 5% BSA in PBS for 1 hour. Primary antibodies against GSDME-N-terminal (Abcam, Cat. ab215191, 1:200), Cleaved CASP3 antibody (CST, Cat. 9664S, 1:200), Cytokeratin 7 (CK7, Abcam, Cat. Ab9021, 1:200), iNOS (Abcam, Cat. Ab3523, 1:200), CD163(Abcam, Cat. ab182422, 1:200), Anti-Mannose Receptor (CD206, Abcam, Cat. ab64693, 1:200) were incubated at 4°C overnight, and sections were washed in PBS (three times, 10 minutes each), followed by incubation with Alexa Fluor secondary antibodies and DAPI (1μg/ml, Sigma, Cat. D9542), as indicated. Washed slides were mounted with Antifade Mountant (Vector, Cat. H1200). Images were taken using a fluorescence microscope (Nikon, Cat. A1R+N-STORM).

### Assembloid whole mount immunofluorescence

Live assembloid in culture were directly fixed in 4% paraformaldehyde (PFA) for 30 min, followed with 60 min of per meabilization and blocking in PBS supplemented with 0.2% Triton X-100 and 5% Bovine Serum Albumin (BSA). For immunofluorescence, assembloid were stained with primary antibodies at 4℃ overnight and sections were washed in PBS (three times, 10 minutes each), followed by incubation with Alexa Fluor secondary antibodies and DAPI (1μg/ml, Sigma, Cat. D9542), as indicated. Washed slides were mounted with Fluoromount-G (Southbiotech, Cat. 0100-01). Images were taken using a Laser Scanning Confocal Microscope (ZEISS 800).

### Flow cytometry

For flow cytometry analysis, single-cell suspensions were obtained by passage through a strainer (70 µm), washed in FACS buffer (PBS with 2% FBS) and stained using the FITC Annexin V Apoptosis Detection Kit Annexin V-FITC/PI (BD, Cat. 556547) or DAPI (Sigma, Cat. D9542, 0.5 μg/ml) according manufacture’s instruction. Flow cytometry was performed on a Thermo Fisher Attune NxT flow cytometer and data were processed with FlowJo software.

### Cell Cytotoxicity and Cell viability assay

The supernatant of trophoblast cells of indicated lines was collected and LDH release was quantified using CytoTox96® Non-Radioactive Cytotoxicity Assay (Promega, Cat. G1780) according to the manufacturer’s instructions. Percentage cytotoxicity was calculated based on maximum LDH release from unstimulated cells lysed with Lysis Solution. Cell viability under 24 hours treatment was determined via the Cell Counting Kit-8 (CCK-8, DOJINDO, Cat. CK04).

### Detection of inflammatory cytokines

Detection of inflammatory cytokines was performed using the LEGENDplex™ Multi-Analyte Flow Assay Kit (Human Inflammation Panel 1 (13-plex), Biolegend, Cat. 740808) following the manufacturer’s instructions. In brief, supernatants of HTR8 were collected after treatment for 24 hours, then diluted 1:1 with assay buffer. 25 μL mixed beads were added to each well and incubated for 2 hours at room temperature with 1000 rpm shaking. The wells were washed once with 1xWash Buffer, followed by detection antibodies incubation for 1 hour, the 25 µL SA-PE was added to each well directly and incubated for 30 min. The beads were washed once and resuspended with 300 µL of 1xWash Buffer. The data were collected on a Attune NxT Flow cytometer (Invitrogen).

### IL-1β, IL-18, BAX and CASP3 ELISA

To measure IL-1β and IL-18 released in vitro, HTR8, TSC or Bewo cells were stimulated with T/S for indicated hours and the supernatants were collected and detected using Duoset ELISA assay kits for human IL-1β (Proteintech, Cat. KE00021) and IL-18 (abcam, Cat. ab215539) according to manufacturer’s protocols. To measure mouse serum IL-1β and IL-18 released in vivo, pregnant mice treated with 0.9% Nacl or T/S were sacrificed, and whole blood was collected at E11.5, followed by centrifuge at 2000g, the serum (50 μl) was extracted and detected with mouse IL-1β ELISA Kit (Cusabio, Cat. CSB-E08054m) and mouse IL-18 ELISA Kit (eBiosciences, Cat. E-EL-M0730c) following by the manufacturer’s instructions. To measure human serum IL-1β, IL-18 and CASP3, the serum (50 μl) was extracted and detected with human human IL-1β (Proteintech, Cat. KE00021), IL-18 (abcam, Cat. ab215539), BAX (GILED BIOTECHNOLOGY, Cat. J20366) and CASP3 (GILED BIOTECHNOLOGY, Cat. J21110) following by the manufacturer’s instructions.

### Statistics and reproducibility

Statistical analysis was performed with GraphPad Prism (v9). Statistical analyses were performed using two-tailed unpaired Student’s t-test and among more than two groups by one-way analysis of variance. Each experiment was performed in three times independently, otherwise indicated. Data were shown as the mean ± SEM, ns, not significant, *P < 0.05, **P < 0.01, and ***P < 0.001, **** P < 0.001.

## Acknowledgement

We thank Prof. Feng Shao from National Institute of Biological Science for kindly share the *Gsdme^-/-^* mice, and Dr. Jia Wang from BeiGene Co., Ltd for data analysis. Work on this project was supported by grants from the National Key Research and Development Program of China (2022YFC2704500 to D.C., 2022YFC2702501 to S.Z. and H.W.), the Key Program of National Natural Science Foundation of China (81830045 to D.C.), the National Natural Science Foundation of China (82171666 to D.C., 82288102 to H.W., 82201861 to S.B.), the Mobility program of Sino German Center (M-0586 to S.Z. and J.K.), the Science and Technology Program of Guangzhou (202201020573 to S.Z.), Guangdong Provincial Basic and Applied Basic Research Foundation (2023A1515110989 to L.Z.).

## Author contributions

B.H., S.B. and Z.F. conducted the experiments; W.D., L.H. and L.Z. collected human samples; Y.W., T.L., L.D., Z.T. and J.K. analyzed the proteomics data. H.W., J. C., D.C. and S.Z. designed experiments and wrote the manuscript.

---

**Supplementary Figure 1.**
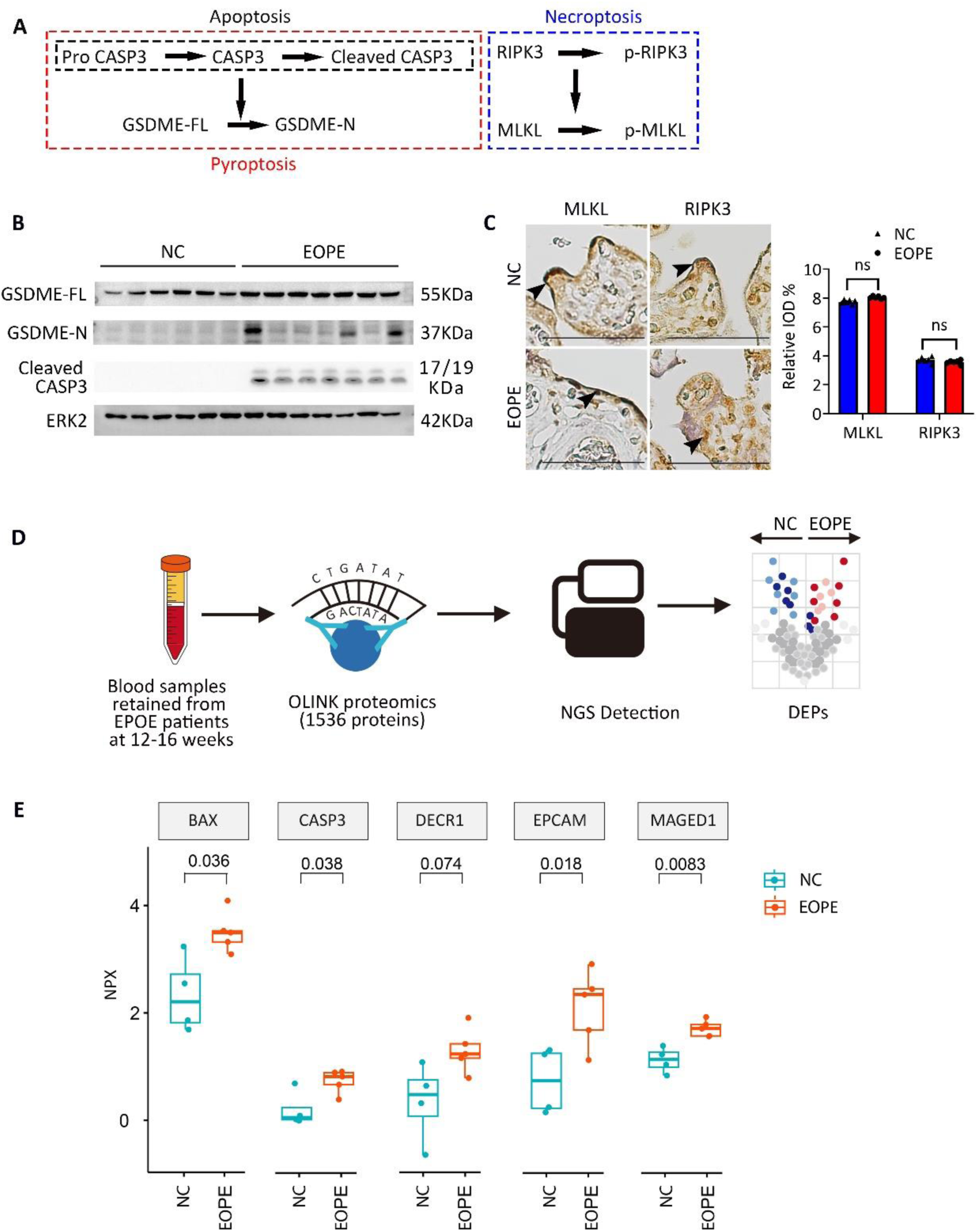
Detection of cell death pathway-related proteins in placental tissues and plasma from EOPE. (A) Schematic diagram of apoptosis, necroptosis and pyroptosis pathways. (B) Immunoblots of GSDME-FL, GSDME-N, and cleaved CASP3 in placental villous lysates from normal controls (n = 6) and EOPE patients (n = 7). ERK2 was used as a loading control. (C) Representative immunohistochemical staining of MLKL and RIPK3 in placental villi from NC and EOPE groups. Arrows indicate positive signals. Similar results were observed in three independent experiments. Scale bars: 50 μm. Relative quantification of MLKL and RIPK3 immunostaining signals in NC and EOPE groups. Error bars represent mean ± SEM. Data were analyzed by Student’s *t*-test; n ≥ 3; ns, not significant.(D) Workflow of blood proteomics using Olink sequencing platform. (E) Box plot showing the expression levels of the five most upregulated proteins among the top 20 differentially expressed proteins.

**Supplementary Figure 2.**
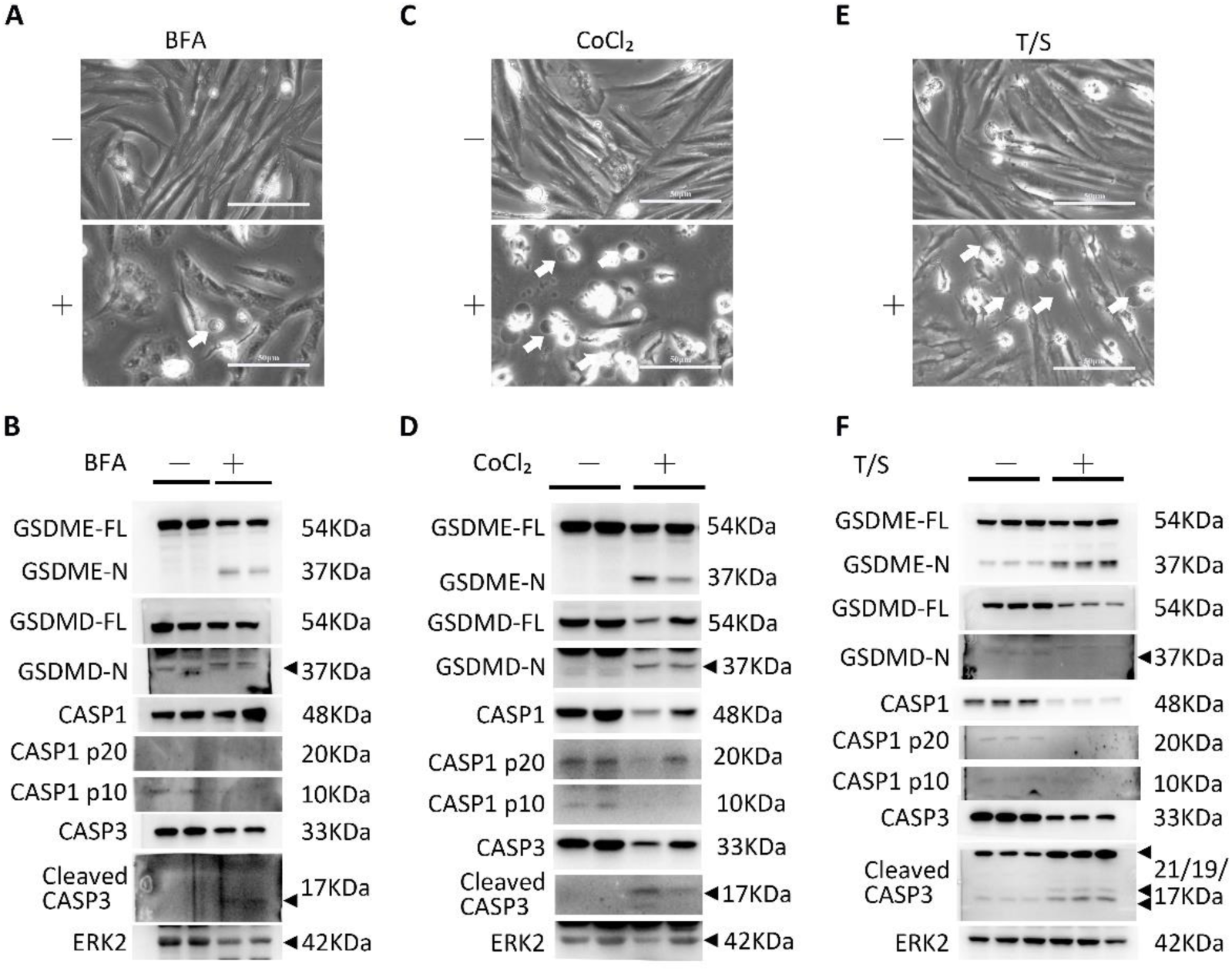
GSDME-mediate pyroptosis was induced upon activation of apoptosis in primary trophoblast cells. (A) Phase-contrast images of primary trophoblast cells (PTCs) treated with brefeldin A (BFA, ER stress inducers), 1μg/ml, after 24 hours. White arrows indicated the pyroptotic-like cells. Scale bars, 50 μm. (B) Immunoblots of GSDME-FL, GSDME-N, GSDMD-FL, GSDMD-N, CASP1, CASP1 p20, CASP1 p10, CASP3 and Cleaved CASP3 in PTCs treated with BFA after 24 hours. ERK2 was used as a loading control. (C) Phase-contrast images of PTCs treated with CoCl2 after 24 hours. White arrows indicated the pyroptotic-like cells. Scale bars, 50 μm. (D) Immunoblots of GSDME-FL, GSDME-N, GSDMD-FL, GSDMD-N, CASP1, CASP1 p20, CASP1 p10, CASP3 and Cleaved CASP3 in PTCs treated with CoCl_2_ after 24 hours. ERK2 was used as a loading control. (E) Phase-contrast images of PTCs treated with SM164 and TNFα (T/S here after) after 24 hours. White arrows indicated the pyroptotic-like cells. Scale bars, 50 μm. (F) Immunoblots of GSDME-FL, GSDME-N, GSDMD-FL, GSDMD-N, CASP1, CASP1 p20, CASP1 p10, CASP3 and Cleaved CASP3 in PTCs treated with T/S after 24 hours. ERK2 was used as a loading control.

**Supplementary Figure 3.**
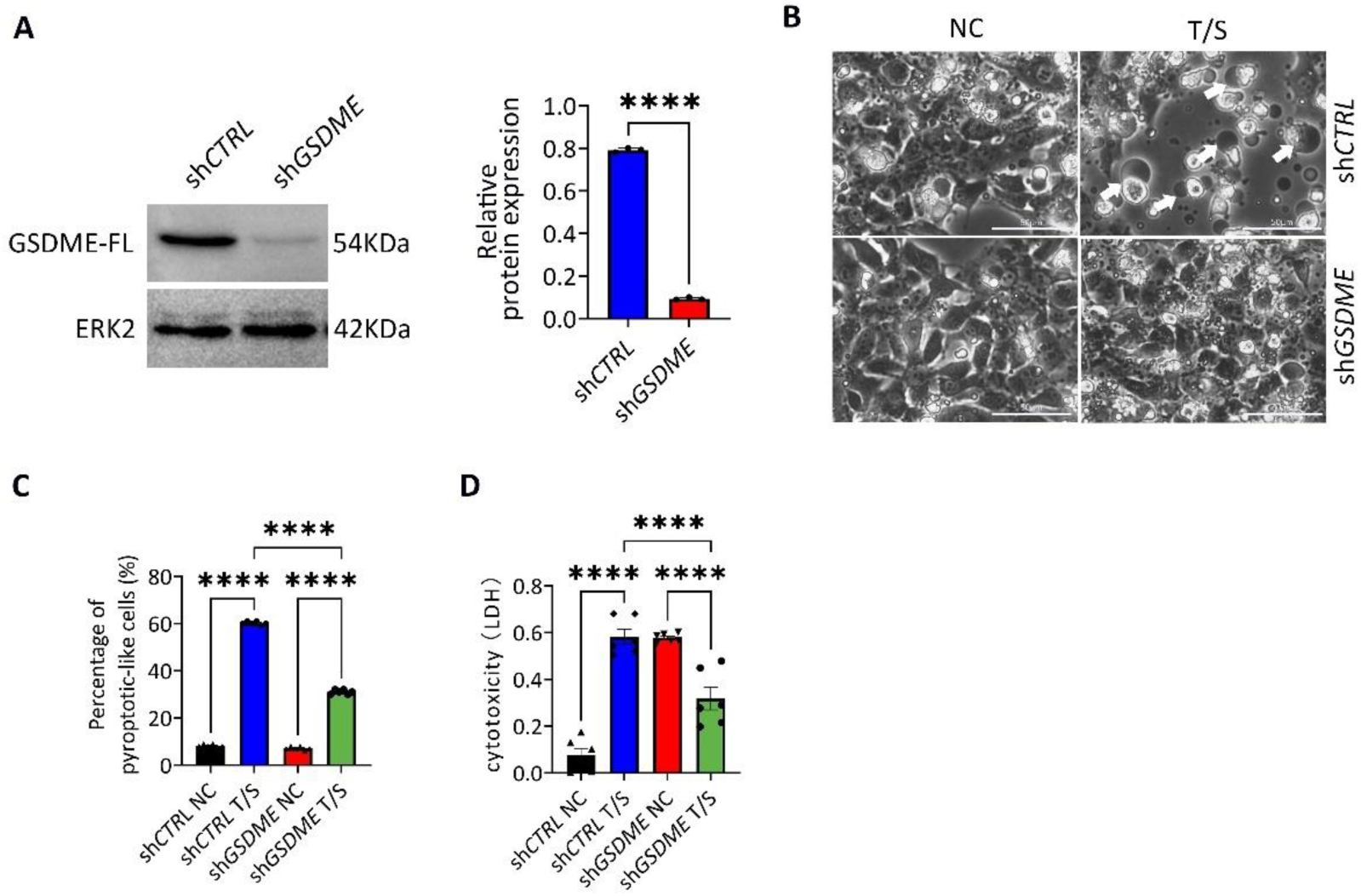
Inhibition of GSDME reduces pyroptosis-like phenotype. (A) Immunoblots of GSDME after shRNA mediated knockdown and its relative quantification (sh*CTRL* and sh*GSDME*) in BeWo cells. ****p<0.0001. (B-C) Phase-contrast images of BeWo cells (sh*CTRL* and *shGSDME*) treated with T/S after 24 hours. Arrows, the pyroptotic cells. Scale bars, 50 μm. (C) Percentages of pyroptotic like cells after T/S treatment. Error bars, mean ± SEM, n ≥ 3. The data were analyzed with a one-way ANOVA. **** p<0.0001. ns, not significant. (D) Comparison of LDH release-based cell death of BeWo cells (sh*CTRL* and sh*GSDME*) treated with T/S after 24 hours. Error bars, mean ± SEM, n ≥ 3. The data were analyzed with a one-way ANOVA, **** p<0.0001, ns, not significant.

**Supplementary Figure 4.**
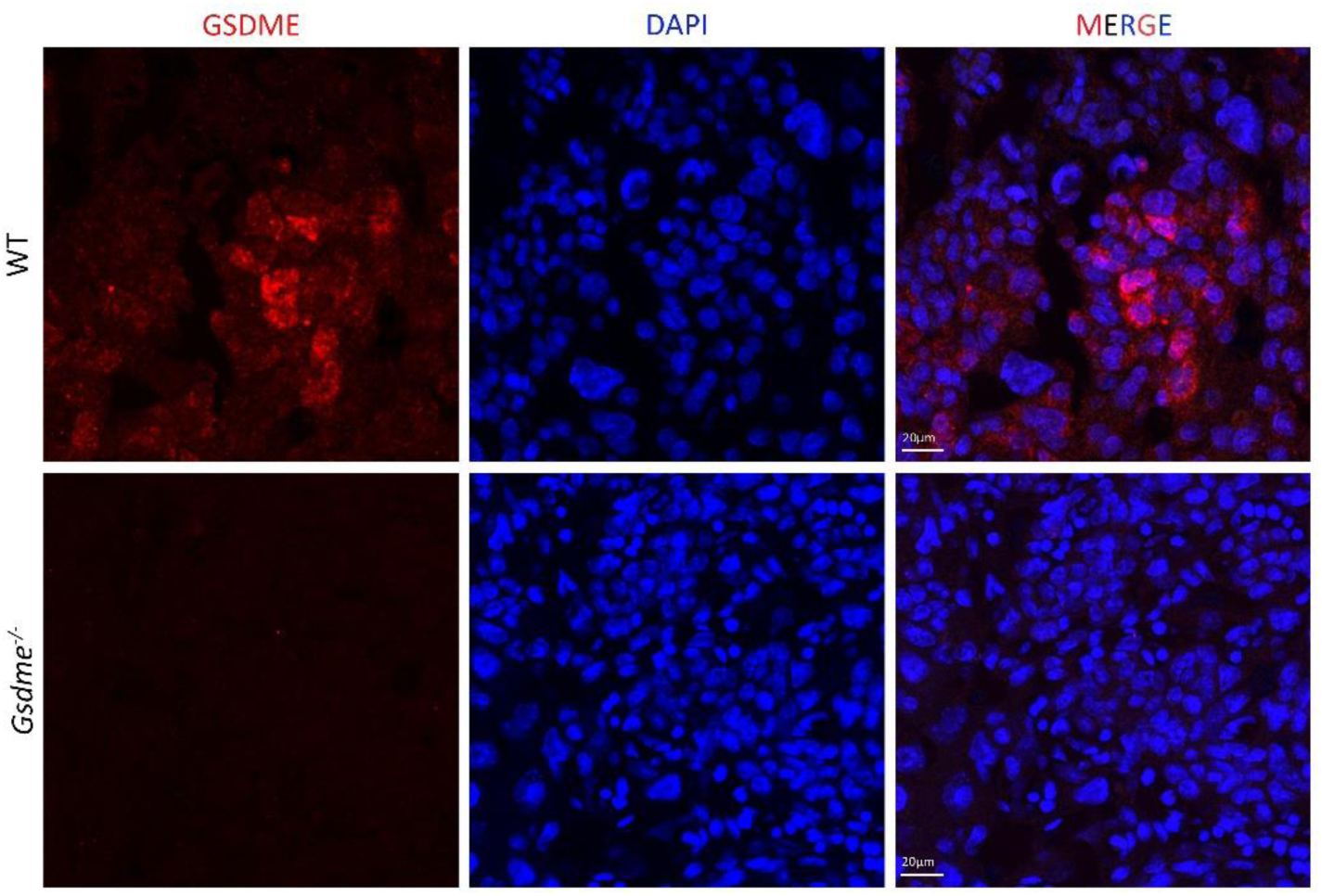
Representative Immunofluorescence staining images of GSDME in placenta tissue of wild type and *Gsdme^-/-^* mice.

**Supplementary Table 1.** Clinical Features of patients for OLINK proteomics.

| group | Sample number | Maternal age(years) | Gestational age at delivery(weeks) | BMI(kg/m<sup>2</sup>) | Primipara | Maximum SBP(mmHg) | Maximum DBP(mmHg) | Neonatal birth weight(g) | Sample collection time(weeks) |
| --- | --- | --- | --- | --- | --- | --- | --- | --- | --- |
| PE | 19B6909223 | 25 | 36.6 | 28.34 | Yes | 150 | 79 | 3040 | 12.9 |
| PE | 19B6909252 | 37 | 34.7 | 31.2 | Yes | 134 | 84 | 2000 | 14 |
| PE | 19B3757678 | 33 | 31.6 | 18.8 | no | 153 | 96 | 1280 | 12.3 |
| PE | 19B2953258 | 36 | 33.9 | 23.53 | no | 128 | 89 | 1670 | 13.9 |
| PE | 19B2084329 | 34 | 27.9 | 26.3 | Yes | 130 | 90 | 670 | 16.4 |
| NC | 19B3983464 | 36 | 37.1 | 22.65 | no | 120 | 75 | 3060 | 12.4 |
| NC | 19B3983179 | 35 | 38.4 | 21.83 | no | 132 | 79 | 2790 | 18 |
| NC | 19B3982793 | 28 | 39.3 | 22.5 | no | 107 | 71 | 2710 | 12.3 |
| NC | 19B2084520 | 36 | 39.7 | 21.2 | no | 122 | 75 | 3260 | 13.6 |

In the control group, preterm delivery was caused by factors such as cervical insufficiency, iatrogenic factors, and preterm premature rupture of membranes.

**Supplementary Table 2.** Clinical characteristics of study participants.

Blood samples for ELISA detection in Figure 1G-H
|  | PE group | control group | P |
| --- | --- | --- | --- |
| sample size | N=11 | N=12 |  |
| Maternal age(years) | 29.83±1.014 | 30.58±1.28 | 0.6508 |
| Gestational age at delivery(weeks) | 38.7 (37.38, 40.0) | 39.65(38.48, 40.08) | 0.1754 |
| primipara, n (%) | 5(41.7) | 6(50) |  |
| Maximum SBP (mmHg) | 146.8±6.804 | 118.0±2.480 | < 0.001*** |
| Maximum DBP (mmHg) | 94.67±3.805 | 76.25±2.918 | < 0.001*** |
| Neonatal birth weight(g) | 2214±104.2 | 2888±83.29 | < 0.0001**** |
| Sample collection time(weeks) | 24.5±0.1508 | 24.42±0.1486 | 0.6977 |

|  | PE group | control group | P |
| --- | --- | --- | --- |
| sample size | N=7 | N=6 |  |
| Maternal age(years) | 31.57±0.7190 | 29.5±0.7638 | 0.0743 |
| Gestational age at delivery(weeks) | 30(29,31) | 31(28.75,32.25) | 0.5541 |
| primipara, n(%) | 3(42.8) | 2(33.3) |  |
| Maximum SBP(mmHg) | 150.4±4.058 | 121±2.978 | < 0.001*** |
| Maximum DBP(mmHg) | 91± | 72.17± | < 0.0001**** |
| Neonatal birth weight(g) | 2100(1540,2500) | 2700(2388,2925) | 0.0261* |

In the control group, preterm delivery was caused by factors such as cervical insufficiency, iatrogenic factors, and preterm premature rupture of membranes.

**Supplementary Table 3.**
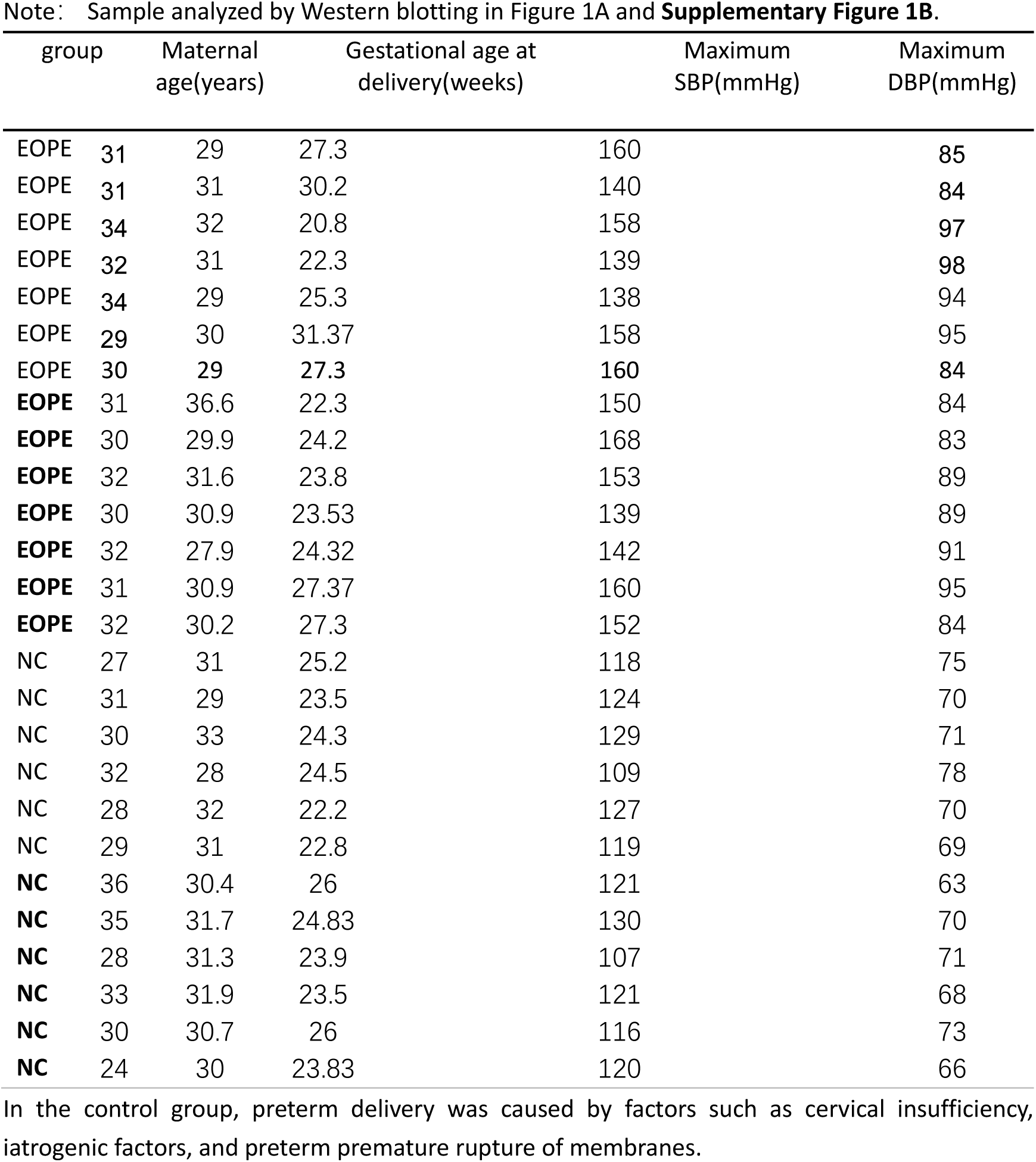
Clinical characteristics of early-onset preeclampsia (EOPE) patients and matched controls.

